# Aging and the encoding of event changes: The role of neural activity pattern reinstatement

**DOI:** 10.1101/809806

**Authors:** David Stawarczyk, Christopher N. Wahlheim, Joset A. Etzel, Abraham Z. Snyder, Jeffrey M. Zacks

## Abstract

When encountering unexpected event changes, memories of relevant past experiences must be updated to form new representations. Current models of memory updating propose that people must first generate memory-based predictions to detect and register that features of the environment have changed, then encode the new event features and integrate them with relevant memories of past experiences to form configural memory representations. Each of these steps may be impaired in older adults. Using functional MRI, we investigated these mechanisms in healthy young and older adults. In the scanner, participants first watched a movie depicting everyday activities in a day of an actor’s life. They next watched a second nearly identical movie in which some scenes ended differently. Crucially, before watching the last part of each activity, the second movie stopped, and participants were asked to mentally replay how the activity previously ended. Three days later, participants were asked to recall the activities. Neural activity pattern reinstatement in medial temporal lobe (MTL) during the replay phase of the second movie was associated with detecting changes and with better memory for the original activity features. Reinstatements in posterior medial cortex (PMC) additionally predicted better memory for changed features. Compared to young adults, older adults showed a reduced ability to detect and remember changes, and weaker associations between reinstatement and memory performance. These findings suggest that PMC and MTL contribute to change processing by reinstating previous event features, and that older adults are less able to use reinstatement to update memory for changed features.

---

Why do humans and other animals remember? One important reason is that features of past experiences can guide current behavior. Recent proposals suggest that a critical function of event memory (1)—also referred to as episodic memory (2)—is to guide anticipation of upcoming events (3, 4). In most cases, using event representations of past experiences facilitates predictions in similar new situations. However, when events unexpectedly change, memory-based predictions are subject to errors. Such errors impose a short-term cost but may have long-term benefits for detecting and registering that features of the environment have changed, as well as for encoding the new event features and integrating them with relevant memories of past experiences to form configural memory representations (5). Thus, memory systems must update representations when things change.

Memory updating upon change detection has been found to depend crucially on interactions between the hippocampus, the surrounding medial temporal lobes (MTL), and the rest of the cortex (6, 7). Memory updating comprises several computational operations with different neural correlates and behavioral signatures (8, 9). These include *pattern completion*, which is the prediction function that activates relevant prior memories and knowledge based on environmental cues, *pattern separation* and *differentiation*, which keep features of the two experiences separate, and *integration*, which captures the relationships between different features of similar events. In order to integrate memory representations of events that are similar but include discrepant features, the brain needs to register the discrepancy and use it to prompt new learning. Models of memory updating propose that when things change, pattern completion leads to prediction errors that can drive new learning, including integration processes to form configural memory representations (5, 10).

These accounts have been supported by behavioral and neuroimaging studies of the learning of word pairs and sequences of words or pictures (8, 9, 11, 12). However, compared to simple laboratory materials, real-world memory updating depends crucially on additional constraints and demands (13). Naturalistic event comprehension relies on a large set of processes working in concert, including object recognition, interpretation of biological motion, spatial orienting, and theory of mind. Event comprehension is also constrained by specific knowledge about particular classes of events and how the world works. For example, when eating a banana, one peels it before eating it. Thus, natural events exhibit correlations across features and time that are more complex than stimuli conventionally used in laboratory settings. Memory systems that capitalize on this richer structure can predict more effectively, but prediction errors and updating of naturalistic activity may function quite differently than the updating of stimuli with simpler temporal and correlational structures. It is therefore important to characterize memory updating in the context of complex, naturalistic activity.

Memory-based prediction and updating may be selectively impaired in older adults. Compared to young adults, older adults are less able to use episodic memory to make explicit judgments about previous events and to predictively guide action (14), and they are particularly prone to error when confronted with events that include overlapping features (15). Such patterns could result from changes to any of several components of memory updating. Behavioral experiments using movie stimuli suggest that when older adults encounter an event that begins similarly to a previous event but ends differently, they are less able than young adults to perform the memory updating necessary for effective formation of configural memory representations (5). In these studies, older and young adults watched movies that included pairs of events that began identically but could end in one of two ways. For example, the actor might unroll a yoga mat and then perform either stretches or abdominal crunches. For both young and older viewers, the ability to detect change, and to remember it later along with the original activity feature, was associated with better memory for the changed features. Older adults detected and remembered fewer event changes, and this was associated with greater memory disruption when a change occurred. These results suggest that, when change is experienced, prediction based on episodic retrieval can drive new learning through the formation of configural memory representations. These results further suggest that this mechanism is less functional in older adults, but the behavioral data alone leave uncertainty about where this breakdown occurs.

Neuroimaging data indicate that patterns of brain activity present while encoding new information are reinstated when this information is recollected, both for simple laboratory materials (e.g., 16, 17) and for more complex stimuli such as movies of everyday activities (18, 19). This effect is usually the strongest in the posterior areas of the Default Network (DN; 20), more specifically part of the posteromedial cortex (PMC) that includes the posterior cingulate cortex (PCC) and retrosplenial cortex (Rsp), and in the medial temporal lobe (MTL) including the parahippocampal cortex (PHC) and hippocampus. These regions are sometimes referred to as the posterior medial system (21) or contextual association network (22) due to their strong involvement in long-term memory recollection, particularly when episodic representations of everyday events must be remembered from visual cues (13, 23).

The hippocampus shows large metabolic alterations and volume loss in aging (24), making functional change in the MTL a potential source of age-related differences in the episodic memory processes that enable the formation of configural memory representations. In addition, the PMC undergoes substantial metabolic and structural change in aging (25), with the integrity of its functioning related to better cognitive abilities in older adults (26). These considerations make the MTL and PMC strong candidates for supporting the reinstatement of event features when encountering a new event that is similar to a previous one. However, there is no evidence to date establishing whether the reinstatement of brain activity patterns facilitates the processing of changes in naturalistic events, nor is there evidence regarding how pattern reinstatement in these regions differs between older and young adults.

In the current study, we aimed to directly assess the role of retrieving episode-specific event features when encoding a new event that was similar to an earlier event. To do so, we used functional MRI (fMRI) in combination with representational similarity analysis (RSA; 27) to assess whether the reinstatement of brain activity patterns associated with past events can facilitate the processing of changes during the perception of new events in older and young adults. We used a task adapted from (5). During fMRI scanning, healthy young and older adult participants viewed two movies of discrete everyday activities, described as two days of an actor’s life (hereafter referred to as Day 1 and Day 2). Together, the activities formed a narrative of the actor’s day. Each activity was made up of two segments: an initial “cue” segment that was always the same on Day 1 and Day 2, and an ending “post-divergence” segment that either repeated or changed on Day 2 (see **Figure 1**). The Day 1 movie consisted of 45 activities. The Day 2 movie depicted activities that were either repeated exactly (15 activities) or began the same but ended differently (30 activities). We stopped each Day 2 movie after the initial cue segment (i.e., before any change) and asked participants to mentally replay the activity ending seen in the Day 1 movie. After this “reinstatement” phase, participants viewed the ending of the Day 2 activity. When the Day 2 movie stopped after each activity ending, participants were asked if they remembered what happened in the Day 1 movie and whether the Day 2 activity ending included a repeated or changed feature (a measure of change detection). We then used RSA to determine the similarity of brain activity patterns in PMC and MTL between the Day 1 viewing and Day 2 reinstatement of activity endings by computing a reinstatement score for each activity and participant.

**Figure 1.**
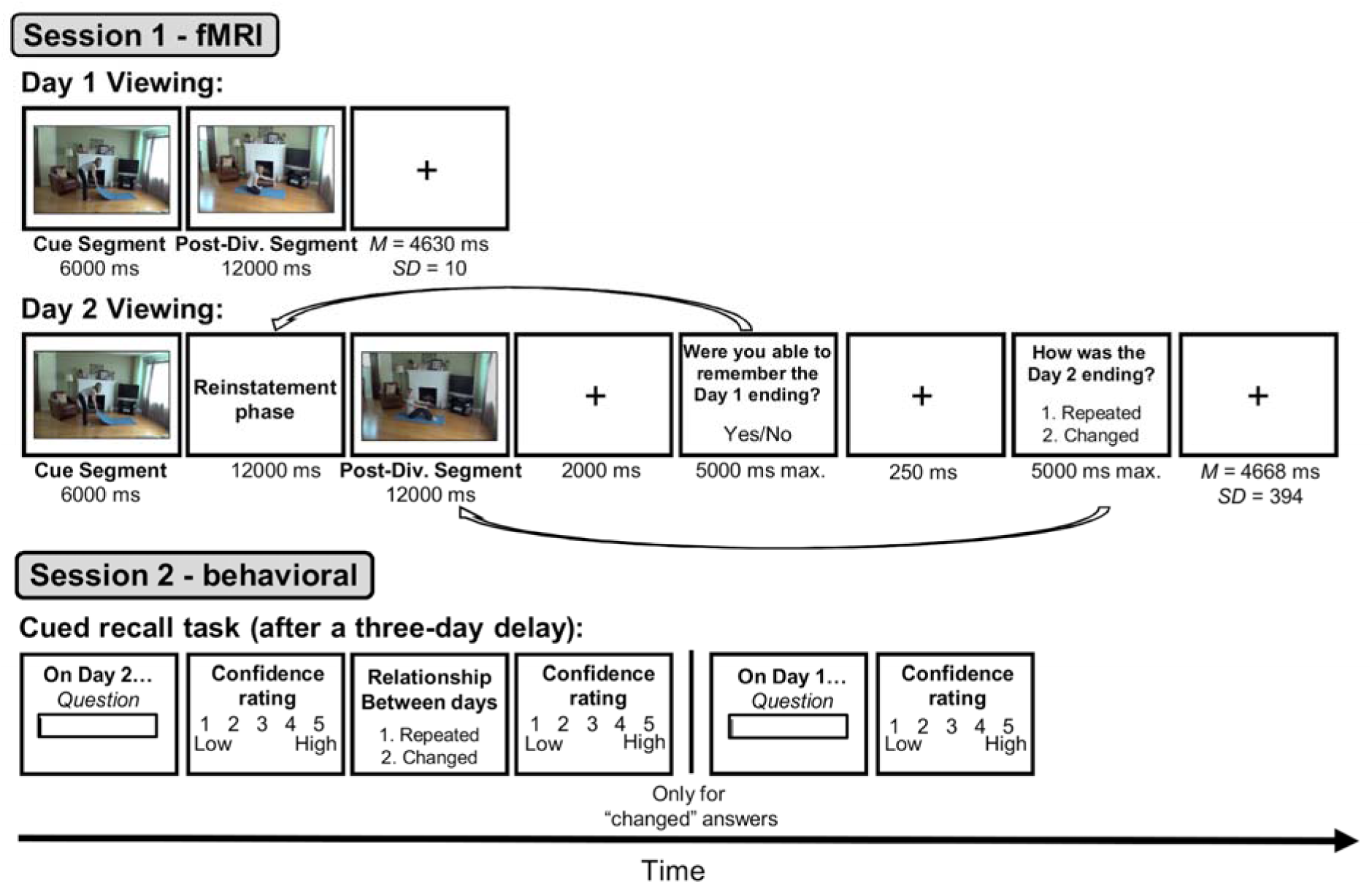
Trial structure of the tasks. For a more detailed description of the materials and procedure, see “Materials, Design, and Procedure,” in the main manuscript and the SI appendix, Section 1.2 and 1.3. Post-Div. Segment = Post-divergence segment.

Three days later, participants were given an unscanned cued recall test for the features of activities viewed in the Day 2 movie (see **Figure 1**). In this memory test, participants were first asked about a feature of the activity (e.g., “What did the actor do on the exercise mat?”), and then were asked whether that activity repeated or changed from Day 1 and Day 2. If they reported that the activity had changed, they were asked to recall the Day 1 feature (see SI, section 1.9. for information on recall scoring).

We hypothesized that stronger Day 2 reinstatement of Day 1 MTL and PMC activity patterns would be associated with better change detection, better subsequent recall of the changed features, and better subsequent recollection that a change had occurred (including what the changed feature had been in the Day 1 movie). Furthermore, given the decline of MTL and PMC functional integrity with aging, we expected that these associations between reinstatement and change detection, memory for changed activity features, and memory for change itself would be stronger for young than older adults and that this difference might be partly explained diminished ability in older adults to retrieve event memories in the service of creating configural memory representations.

## Results

All analyses of memory performance and neural pattern reinstatement effects were computed using linear mixed effects models with subjects and activities as random effects. Logistic models were used when the dependent variable was binary (see the SI appendix, Section 1.8 for more details).

### Memory performance and change classifications

During Day 2 viewing in the scanner, participants were asked after each activity whether they had successfully reinstated the Day 1 activity ending and whether its Day 2 ending was the same as on Day 1 or had changed (**Figure 1**). The reported rates of reinstatement success were higher for older than young adults, but older adults were less accurate at detecting when activities had changed. In young but not older adults, self-reported reinstatement predicted change detection accuracy. These results suggest that older adults were overconfident compared to young adults and less able to use subjective features of reinstatement to detect changes (see SI Appendix section 2.1.1 for a detailed statistical report).

We examined performance in the unscanned cued recall task to determine whether remembering change and the original Day 1 activity was associated with better memory for the changed Day 2 features, and whether this was affected by age. Participants attempted to recall event features from the Day 2 movie, and then were asked whether that event had changed from Day 1 to Day 2. When participants indicated that an event had changed, they were asked to recall the Day 1 feature. Change could therefore be remembered with recall of Day 1 features (change recollected), remembered without recall of Day 1 features (change remembered, but not recollected), or not remembered at all.

Models including fixed effects of age group and activity type indicated that older adults recalled fewer Day 2 features than young adults [χ*^2^*(1) = 9.61, *p* = .002], and that this effect of age did not differ between repeated and changed activity types [χ*^2^*(1) = 0.37, *p* = .54] (see **Figure 2**). To examine the association between change recollection and Day 2 recall, we fitted another model with fixed effects of age group and activity type but with levels for changed activities corresponding to each type of memory for change. Both age groups recalled changed activities less accurately than repeated activities when change was not remembered at all or remembered, but not recollected (*smallest z* ratio = −7.64, *p* < .001). However, when participants recollected change, recall was higher than for repeated activities (*z* ratio = 5.95, *p* < .001). The estimated probabilities of change recollection were lower for older (*b* = .27, *95% CI* = [.19, .36]) than young (*b* = .38, *95% CI* = [.29, .48]) adults, [χ^2^ (1) = 4.70, *p* = .03]. Thus, both age groups showed enhanced recall of Day 2 features when change was recollected, but older adults experienced this benefit on fewer trials.

**Figure 2.**
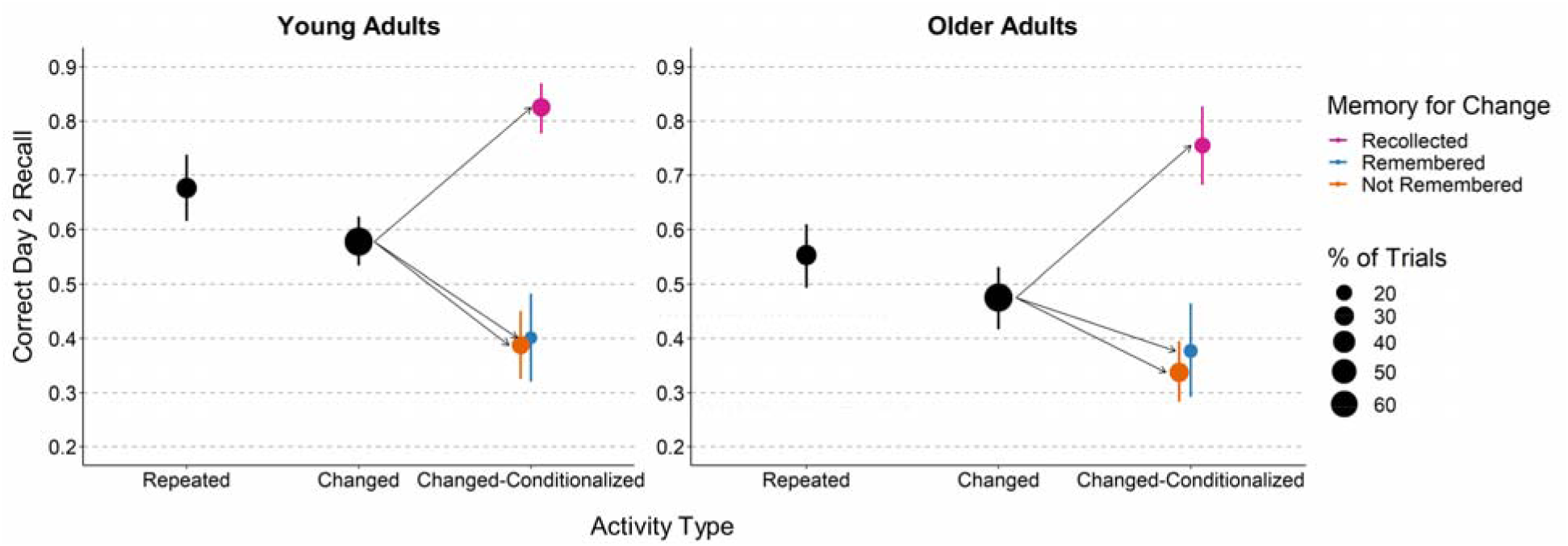
Mean probabilities of correct Day 2 recall. Error bars are bootstrap 95% confidence intervals. Change could be remembered with recall of Day 1 features (Recollected; purple points), remembered without recall of Day 1 features (Remembered; blue points), or not remembered at all (Not Remembered; orange points). Point areas for conditional cells reflect the relative proportions of observations in each cell.

For both young and older adults, self-reported successful reinstatement of Day 1 features was associated with better recall of the changed Day 2 features. However, this effect was not significant after controlling for variation in change recollection, indicating that self-reported successful reinstatements predicted better recall of changed Day 2 features because those reinstatements also predicted better memory for change and recall of the Day 1 activity feature at test (see SI appendix, Section 2.1.1). When participants could not recall changed activity features, they were likely to erroneously intrude features from the corresponding activities viewed on Day 1. Analyses of such Day 1 intrusion rates generally mirrored the rates of correct Day 2 recall (see SI appendix, Section 2.1.2).

### Representational similarity measures of neural memory reinstatement

To assay the neural reinstatement of activity-specific Day 1 features following cue segments on Day 2, we compared neural activity patterns in the MTL and PMC during Day 1 viewing to the patterns from attempted reinstatement of Day 1 activities. Patterns were analyzed at the level of parcels in the 17 networks/300 parcels cortex parcellation map from (28), focusing on PMC and MTL parcels (see **Figure S1**).

To extract patterns for each Day 1 post-divergence segment and Day 2 reinstatement attempt for each participant, we averaged the BOLD signal for each voxel over the ninth to fourteenth scans (11.97 to 18.62 s) after the beginning of the cue segment for each activity; this interval encompasses the fMRI response to the Day 1 post-divergence segment / Day 2 reinstatement phase, accounting for shift due to hemodynamic lag. We then computed reinstatement Z-scores that quantified the degree to which reinstatement activation patterns are more similar to their *matching* Day 1 activity encoding activation pattern than to the others (18, 19; see **Figure 3** for details). Reinstatement Z-scores for each parcel within each ROI were averaged, resulting in two scores for each activity for each participant, one for the MTL and one for the PMC.

**Figure 3.**
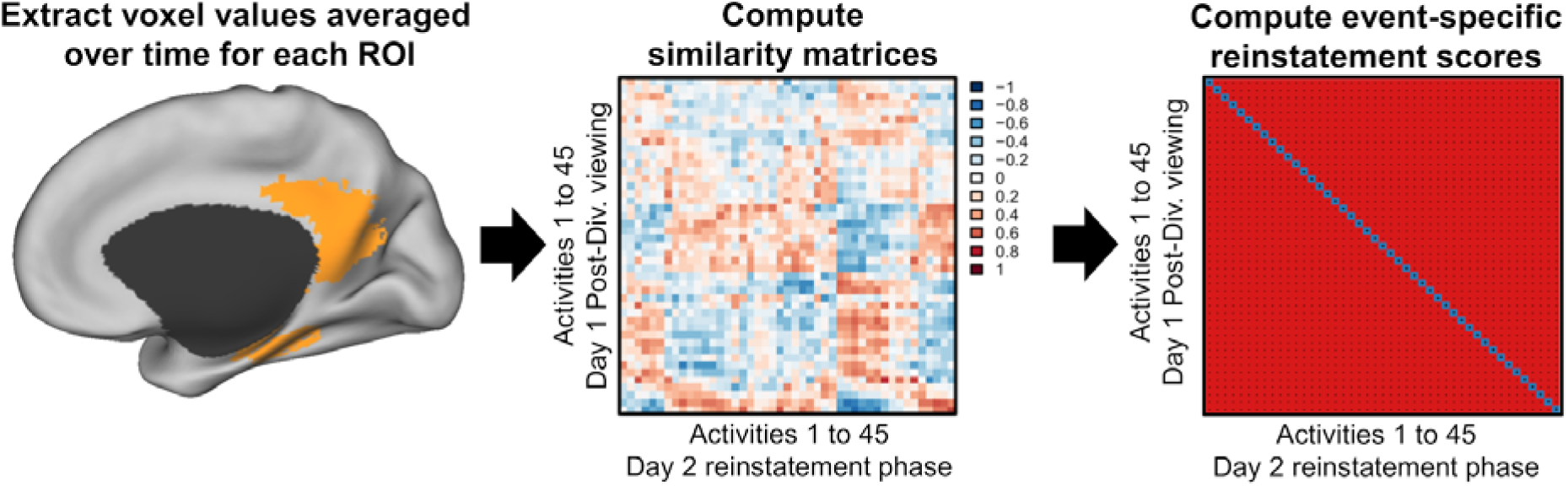
Multivariate voxel analyses. After averaging voxel values over time in the parcels of interest during the Day 1 post-divergence segments and Day 2 reinstatement phases for each activity and participant, we correlated voxel values in the parcels of interest for each activity during Day 1 viewing with the corresponding voxel values for all the activities during Day 2 reinstatement. The resulting correlation coefficients can be plotted in similarity matrices, as illustrated above. We then computed event-specific reinstatement scores by calculating the differences between each of the on-diagonal values indicated in blue and the off-diagonal values indicated in red. To calculate the distribution of this measure under the null hypothesis of no reinstatement, we randomly permuted the labels of the Day 2 activities (i.e., the columns in the middle panel) 1,000 times and recomputed the correlation difference for each random permutation. The final reinstatement Z-score was the ranking of the actual (unpermuted) difference score relative to its null distribution, transformed to a Z-score.

Reinstatement Z-scores were mostly positive, indicating that participants were able to reinstate activity-specific neural activation patterns (see **Figure 4A**). Linear models with no fixed effect and reinstatement Z-scores as dependent variables indicated that the intercept was significantly above zero for both the PMC (*b* = 0.17, *95% CI* = [.12, .23], *t*-value = 6.49, *p* < .001) and MTL (*b* = 0.14, *95% CI* = [.08, .20], *t*-value = 4.77, *p* < .001). Similar models with age group added as a fixed effect revealed no age differences [PMC: χ^2^ (1) = 0.07, *p* = .79; MTL: χ^2^ (1) = 2.01, *p* = .16]. Follow-up analyses on the individual parcels within the cortex parcellation map (28) indicated significant reinstatement effects in many parcels for both age groups (see **Figure S2**), only one parcel within the two ROIs showed a significant age difference in the reinstatement effect (Left MTL parcel 144; see **Table S1**).

**Figure 4.**
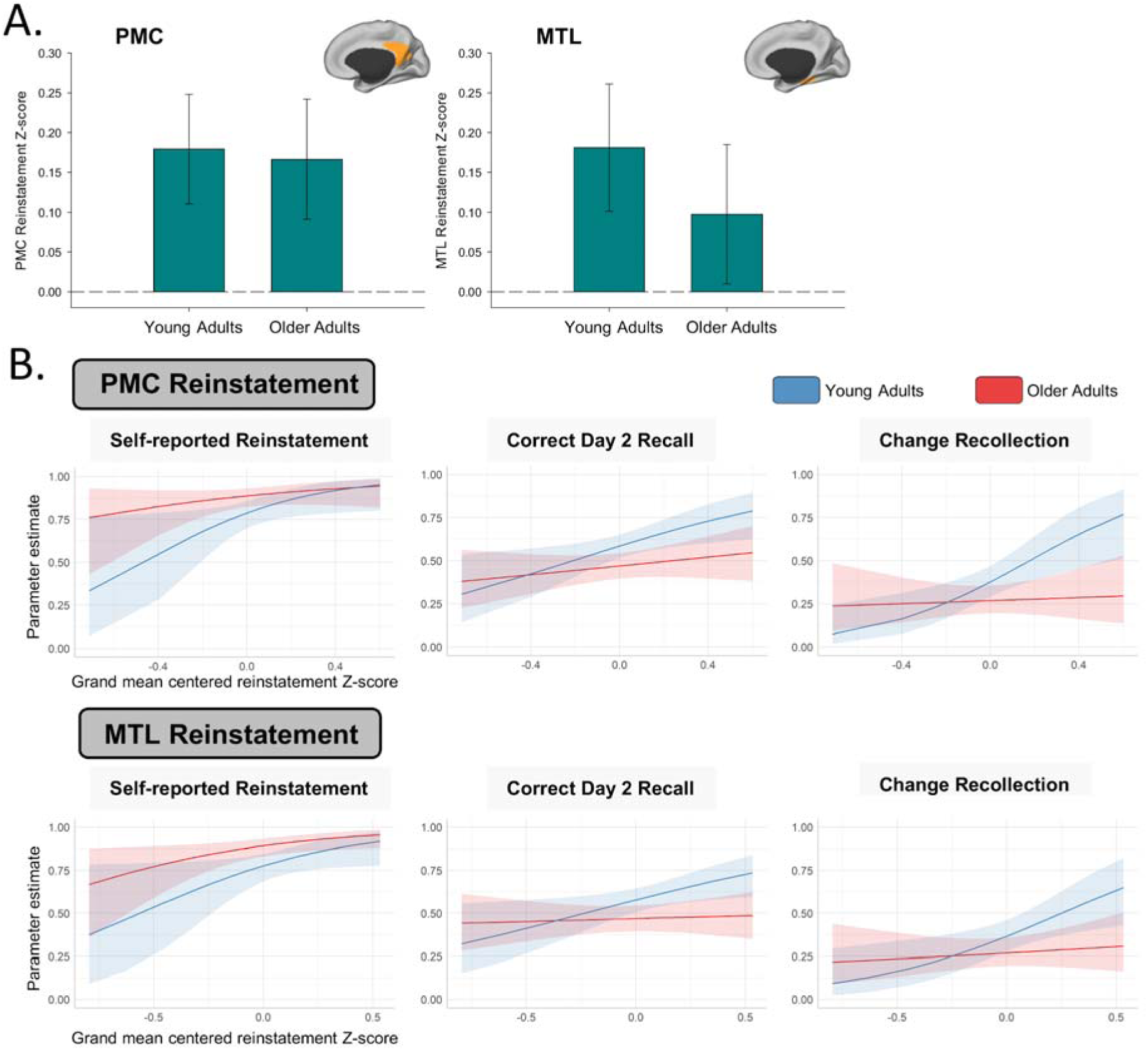
Panel A: Parameter estimates for mean reinstatement Z-scores in the PMC and MTL by age group. The error bars are 95% confidence intervals. Panel B: Parameter estimates for the between-participant associations between mean PMC/MTL reinstatement Z-scores and behavioral memory measures. The shaded regions are 95% confidence intervals.

We then fitted models with between- and within-participant reinstatement Z-scores for changed activities, as well as age group, as fixed effects. Between-participant reinstatement Z-scores were mean reinstatement Z-scores for the changed activities of each participant across all changed activities for that participant. These allowed us to examine whether participants who reinstated Day 1 neural activity patterns more strongly also recall of Day 2 features more accurately. Note that the grand mean (i.e., the mean of the mean reinstatement Z-scores) was subtracted from all observations for plotting. Within-participant reinstatement Z-scores were computed by centering reinstatement Z-scores for the changed activities within each participant (i.e., the mean of reinstatement Z-scores from all changed activities of that participant was subtracted from the reinstatement Z-score of each changed activity). These allowed us to examine whether Day 2 activities with higher reinstatement Z-scores (independent of the mean reinstatement of the participants) were associated with better subsequent memory for Day 2 features. In addition, because reinstatement Z-scores across parcels were only moderately correlated (**Table S2**), we performed similar analyses but examined the effect of each individual PMC and MTL parcel on Day 2 recall performance above and beyond the effects of all the other parcels in the ROI. This procedure allowed us to examine whether reinstatement in specific parts of the PMC or MTL (such as the hippocampus) predicted behavioral performance. For brevity, only summarized results of these latter analyses are presented below (see the SI Appendix, Sections 2.2.2 and 2.2.3 for a more detailed statistical report).

#### Between-participant differences in reinstatement Z-scores were related to memory performance and subjective experience

We first examined whether RSA reinstatement Z-scores were correlated with *self-reported reinstatement success* across participants (**Figure 4B**, left panels). Participants with higher mean PMC reinstatement Z-scores rated more of their neural reinstatements as successful [χ^2^ (1) = 5.44, *p* = .02]. Although this effect did not interact with age group [χ^2^ (1) = 0.89, *p* = .35], it was driven by specific PMC parcels in young but not older adults (see SI Appendix, section 2.2.2). There was also a significant effect of mean reinstatement Z-score for the MTL [χ^2^ (1) = 9.02, *p* = .003] that did not interact with age group [χ^2^ (1) = 0.10, *p* = .75] and was not driven by specific parcels. Further, mean PMC and MTL reinstatement Z-scores were correlated with the accuracy of *change detection judgments* in young but not older adults. Reinstatement in different PMC parcels both positively and negatively predicted change detection in each age group, with more parcels being negatively than positively associated with change detection for older adults (see SI Appendix sections 2.2.1 and 2.2.2), suggesting that older adults were less able to use reinstated activities to detect changed features.

We next correlated the mean reinstatement Z-scores for changed activities with *correct Day 2 recall performance* (**Figure 4B**, middle panels). Participants with higher PMC mean reinstatement Z-score had more accurate recall of Day 2 features [χ^2^ (1) = 5.43, *p* = .02], and this effect did not interact with age group [χ^2^ (1) = 1.85, *p* = .17]. For the MTL, between-participant mean reinstatement Z-scores did not predict Day 2 recall [χ^2^ (1) = 2.62, *p* = .10], and reinstatement Z-scores did not interact with age group either [χ^2^ (1) = 2.78, *p* = .09]. Examining the effects of individuals parcels revealed that these between-participant effects were not driven by specific PMC or MTL parcels. For the analyses of Day 1 intrusions during Day 2 recall of changed activities, see SI Appendix, Section 2.2.1 and 2.2.2.

Third, we examined whether reinstatement Z-scores predicted *change recollection* (**Figure 4B**, right panels). Given that Day 2 recall performance was comparable when change was remembered without Day 1 recall and when change was not remembered at all (see **Figure 2**), we collapsed these cells here and in subsequent analyses into the category *change not recollected*. Consistent with Day 2 recall, mean between-participant PMC reinstatement was positively associated with change recollection [χ^2^ (1) = 4.44, *p* = .04], but the effect was qualified by an interaction with age group [χ^2^ (1) = 4.71, *p* = .03]. There was a significant positive association for young adults [χ^2^ (1) = 8.05, *p* = .005] but not for older adults [χ^2^ (1) = 0.09, *p* = .76]. Analyses of individual parcel Z-scores revealed that the effect was driven by specific parcels in the young adults only (see SI Appendix, section 2.2.2). There was a significant effect of mean reinstatement Z-score in the MTL [χ^2^ (1) = 4.05, *p* = .04] that did not interact with age group [χ^2^ (1) = 3.01, *p* = .08], indicating that participant with higher mean MTL reinstatement Z-cores had a better change recollection accuracy.

Finally, because between-participant differences in mean PMC reinstatement Z-scores for young adults were positively related to both Day 2 recall of changed features and to change recollection, we examined whether the association between reinstatement Z-scores and Day 2 recall accuracy for the changed activities could be explained by change recollection. This model included change recollection accuracy and mean between-participant PMC reinstatement Z-score as fixed effects (see **Figure S3** for the associations between mean reinstatement Z-scores and Day 2 recall, and for the unique contribution of change recollection to Day 2 recall). With both reinstatement Z-scores and change recollection in the model as fixed effects, PMC reinstatement no longer predicted Day 2 recall [χ^2^ (1) = 1.69, *p* = .19]. However, change recollection was still positively associated with Day 2 recall [χ^2^ (1) = 147.40, *p* < .001]. The interaction between reinstatement Z-scores and change recollection accuracy was not significant [χ^2^ (1) = 2.10, *p* = .15]. This is consistent with the possibility that the association between reinstating of Day 1 activities during Day 2 viewing and subsequent Day 2 recall is mediated by recollecting that the activity had changed. However, the current design does not allow for strong causal conclusions about this potential mediation.

#### Within-participant differences in reinstatement Z-scores were related to memory performance

Between-participant analyses informed how individual differences in reinstatement Z-scores were related to behavioral performance. To assay whether activity-to-activity variation in reinstatement within a person was related to subsequent memory, we conducted a second set of analyses. First, Regarding Day 2 recall, neither the effect of mean reinstatement Z-scores in the PMC and MTL nor the interaction with age group was significant [*largest* χ^2^ (1) = 2.23, *p* = .14, see **Figure S4**]. However, analyses of individual parcels revealed that reinstatement Z-scores in two PMC parcels were positively associated with Day 2 recall, whereas one other parcel showed the opposite effect, and no interaction with age group (see SI Appendix, Section 2.2.3). There was no significant effect of individual parcel reinstatement nor was there an interaction with age group in the MTL. No significant effects were found for either the mean PMC or MTL or individual parcel reinstatement Z-scores regarding change recollection, self-reported reinstatement success, change detection accuracy, or Day 1 intrusions onto Day 2 recall [*larges*t χ^2^ (1) = 3.01, *p* = .08], with the exception of the interaction between age group and mean PMC reinstatement Z-score for change detection accuracy [χ^2^ (1) = 4.63, *p* = .03]. Reinstatement Z-scores were not related to change detection accuracy in the young adults [χ^2^ (1) = 0.08, *p* = .77] but greater mean PMC reinstatement predicted poorer change detection in the older adults [χ^2^ (1) = 4.87, *p* = .03]. Thus, apart from Day 2 recall of changed activities, there was no evidence for a beneficial within-participant relationship between neural pattern reinstatement and behavioral memory performance in either age group.

### Mass univariate analyses

To further investigate age differences in the processing of changed Day 2 activities during viewing, we performed mass univariate fMRI analyses using general linear models, as described in the SI appendix, Section 1.6. Many models of memory updating propose that novelty detection and prediction error are critical components of memory updating when encountering changes (5, 7, 10). Consequently, we specifically examined the difference in the neural response between viewing of changed and repeated activity endings (the “post-divergence” segments). Across all participants, we found more activity for changed than repeated endings in clusters mainly located in the lateral prefrontal cortex and, at a lower threshold, in the bilateral anterior hippocampus (see **Figures S5 and S6**). A two-sample *t*-test showed that this neural response did not differ between age groups (no cluster was activated above threshold). In addition, neural activity was not parametrically modulated by reinstatement Z-scores during the changed activity epochs in any brain region for either age group (again, no cluster was activated above threshold).

To further assess the proposal that hippocampal responses to novelty are a critical component of memory updating when experiencing changes in events (e.g., 10), we next examined whether between-participant differences in hippocampal response intensity while viewing changed versus repeated activity endings predicted reinstatement Z-scores and behavioral measures. Results showed no association between hippocampal response intensity and either PMC or MTL reinstatement Z-scores. At the behavioral level, hippocampal responses were only positively associated with change detection during Day 2 viewing. No interaction with age group was significant (see SI Appendix, Section 2.3 for details).

Finally, an unexpected finding of the RSA analyses was that reinstatement Z-scores did not differ across age groups. To examine whether there might still have been age-related differences in how participants initially perceived the activities, we used a pattern classifier which showed that the voxelwise distribution of activation in the PMC and MTL differed between young and older adults (see SI Appendix section 1.7 and 2.4). Thus, although young and older adults did not differ in their neural reinstatement of Day 1 activities during the Day 2 reinstatement phases (see **Figure 4A**), the neural activity patterns during Day 1 viewing still differed between the two groups.

## Discussion

In this study, attempting to reinstate features of a relevant previous event during comprehension of a new one was associated with widespread reinstatement of fMRI activity patterns corresponding to anticipated features of the event. As hypothesized, reinstatement was associated with better subsequent memory for changed event features in the PMC. This was true for the PMC as a whole at the between-participant level, and for a subset of PMC parcels at the within-participant level. When participants attempted to recall changed event features, the ability to recall *what* the feature had changed to was associated with being able to recollect *that* the feature had changed and to report what it had been previously. In addition, the positive association between neural pattern reinstatement in the PMC and memory for changed activity features was statistically explained in the young adults by their ability to recollect the original activity feature and the fact that the activity had changed. A similar pattern was seen for subjective judgments about whether participants had successfully reinstated activity features before viewing each activity’s ending. In the MTL, between-participant differences in reinstatement for the ROI as a whole significantly predicted better change detection and memory for the original activity features; and the intensity of the neural response in the anterior hippocampus while viewing changed events predicted better change detection and recognition performance. However, reinstatement in the MTL was not related to better recall of the changed event features.

Neural measures of pattern reinstatement in the PMC as a whole were associated with more accurate recollection of the new information presented *after* the end of the reinstatement phase, information that *conflicted* with the previously encoded (and reinstated) features. This finding is consistent with previous studies showing that during recollection, reinstatement of the brain activity pattern present in posterior DN areas while watching movies can predict memory for the movie content up to one week later (18, 19), and with evidence for neural pattern reinstatement when rehearsing learned associations to pictures (9). The current results indicate for the first time that such reinstatement is related to the encoding of novel, unexpected event features. One possibility is that reinstatement leads to predictions, which in turn lead to a prediction error signal when events change, and then to memory updating (11). In a previous study using sequences of pictures, updating manifested as selective forgetting, or *pruning,* of previously encoded features (11), whereas here memory updating was associated with better memory for previous as well as new event features. We attribute this difference to the formation of a configural representation composed of the original activity features, the changed features, and their temporal relations (5). Consistent with this idea, both neural and behavioral measures of reinstatement before encoding the change were associated with being able to recollect how the activity had changed, and neither self-reported nor neural pattern reinstatement remained significant predictors of memory accuracy for the changed features after controlling for change recollection accuracy. Forming a configural representation is related to *integration* of separate experiences into a common context, which is also associated with cortical reinstatement (9). It can be contrasted with *differentiation*, in which overlapping features are selectively deleted; differentiation acts to make experiences more distinctive rather than to merge them into a complex (29).

The PMC may play a key a role in supporting the event model representations from which predictions are generated. The PMC is part of the DN, and it was initially thought to be primarily involved in generating internal mentation, which stands in opposition with attention to the external world (30). However, there is now substantial evidence that the PMC also supports externally-directed attention and event comprehension when task performance and the processing of perceptual inputs can benefit from relevant information stored in memory (31). In addition, recent studies have revealed that the transitions between activity patterns within the PMC while watching movies follow timescales ranging from seconds to minutes, closely matching how people segment movie content into distinct events (32). This supports the view that event model representations might be the means by which PMC facilitates interactions with the external world. Interestingly, at the level of individual parcels, reinstatement in some PMC parcels negatively predicted memory accuracy for the changed features, change detection performance, and self-reported reinstatement success. This suggests that the PMC might not be unitary regarding the role of its subregions in cognition, which aligns with recent speculations that some subregions might be more involved in processing perceptual inputs than memory representations (25).

As for the MTL, there is extensive evidence that this region is involved in the relational binding of information stored in memory and how it relates to perceptual inputs in order to form associative memory representations of everyday experiences (33, 34). Consistent with these findings, research has shown that peaks of activity in the hippocampus at the transition between perceived events can predict neural reinstatement in the PMC during recall (32, 35). Pattern reinstatement within the MTL in the current study might therefore reflect the relational binding of information stored in memory—whose retrieval is triggered by the cue segment—in order to form the event model that is supported by the PMC. This fits well with our results showing MTL involvement in detecting change and remembering original event features. However, MTL reinstatement did not predict memory for the changed features, suggesting that the formation of configural memory traces of everyday events relies more strongly on the contribution of cortical areas, among which the PMC might play a prominent role.

Older adults were less likely than their younger counterparts to detect change and to recall the original activity features during the cued recall task. A possible explanation for this finding is that older adults are less able to use retrieved activity features when encoding changed features to form an updated configural representation that includes both features and their relationship. Previous studies of age-related deficits in change comprehension (5) and in associative memory (36) are consistent with this idea. A possible explanation for our findings is suggested by behavioral studies showing that the individuation of events during perception is impaired in older people (37): the event representations formed by older adults during the original viewing of the activities may have been less detailed than those of young adults. The finding that the spatial pattern of activation during movie viewing differed across groups is consistent with this possibility. As a result, they may have been less successful in forming configural representations when confronted with changed activities, and thus less successful in encoding and remembering changes. Although speculative, this proposal might explain why self-reported reinstatement accuracy in older adults did not predict better change detection during Day 2 viewing and was not associated with fewer Day 1 intrusions (See SI Appendix, Section 2.1.2) nor accurate detection of changed activity features on Day 2 (See SI Appendix, Section 3.2), as was the case for young adults. Further studies examining the quality of encoding during the initial viewing, for instance by asking participants to verbalize their experience while watching the Day 1 movie and then relating these verbal reports to neural activity patterns would test this possibility.

In any functional neuroimaging study comparing young and older adults, it is important to consider potential sources of artifact; these include group differences in neurovascular coupling, head motion, and how the tasks are approached (38). Here, the fact that older adults showed robust overall neural reinstatement renders their significantly weaker relationships between neural reinstatement and behavioral memory measures particularly striking. Another caveat was that participants were instructed to reinstate previous activity features while Day 2 activities were paused. Cognitive age differences are often larger when under time pressure and when self-initiated processing is required (39); therefore, the time to deliberately reflect in our study may have attenuated age differences. To generalize to naturalistic comprehension, it will be important to use converging measures that do not depend on strategic, interruptive task instructions.

In conclusion, the present results showed that the reinstatement of previously generated responses in the PMC and MTL predicted better memory for reinstated activity features and change detection, and that PMC reinstatement facilitated the encoding of related but changed activity features. This latter finding is particularly striking because the changed features conflicted with the just-retrieved features of the previous activity. We propose that retrieving activity features facilitates encoding precisely because it enables the system to register discrepancy between the predicted and encountered features. That discrepancy can drive formation of a configural representation that includes the old features, the new ones, and their relationship. This process was impaired in older adults; further, the pattern of impairment suggests that deficits in encoding a detailed memory representation of the original event might reduce older adults’ ability to encode a configural representation of the changed event that includes its relationship to the previous event.

## Method

The full stimulus sets for the materials used in the present experiments, anonymized data files, coded data, and R Markdown files (40) containing the analysis scripts are available on the Open Science Framework: (https://osf.io/v3dqg/).

### Participants

This study was approved by the IRB of Washington University in St. Louis. All participants gave their written informed consent before participating in the study. Participants were recruited from the Washington University School of Medicine Research Participant Registry, flyers posted on campus, and word of mouth. Potential participants were initially contacted by phone for a prescreening interview. The sample included 62 healthy right-handed participants: 34 young adults (mean age 22.85 years, *SD* = 2.71, range: 18-27 years, 22 females) and 28 older adults (mean age 69.86 years, *SD* = 5.01, range: 65-84 years, 20 females). All older adults had a score of 27 or above (*M* = 29.25, *SD* = 0.87, range: 27-30) on the Mini-Mental State Exam (MMSE; 41). For more details about recruitment and exclusion criteria, see the SI Appendix Section 1.1.

### Materials, Design, and Procedure

The materials were movies of a female actor performing daily activities on two fictive days in her life, which were described to participants as “Day 1” and “Day 2” (5). There were 45 activities, each of which was filmed in two versions (A and B) that differed on a thematically central feature (e.g., doing stretching or sit-ups on a yoga mat, see **Figure 1**). Each activity began with a 6,000 ms initial cue segment that was consistent across versions, followed by a 12,000 ms post-divergence segment (A or B). The version of the activity that participants saw in the Day 1 movie (A or B) and whether the activity repeated or changed in the Day 2 movie were both counterbalanced across participants.

Participants viewed the Day 1 and Day 2 movies during fMRI scanning, and then returned after 3 days for the memory tests (see **Figure 1**). Activities in the Day 1 movie appeared as continuous 18,000 ms clips that were each followed by a fixation interval. After a delay of approximately 10 min, during which field map and anatomical images were collected, participants then watched the Day 2 movie. All Day 2 activities were paused for 12,000 ms between the cue and post-divergence segments (repeated or changed), during which participants were asked to mentally replay the Day 1 ending. They were then asked whether they successfully replayed the Day 1 features when the movie was stopped, and whether the activity features had changed. Finally, we collected a second set of field map images and a high-resolution T2-weighted image, taking approximately 6 minutes. During Session 2 (outside the scanner), we first tested participants’ memory for the previously viewed activities using a cued recall task (see **Figure 1**). The recall cues for each activity appeared in the same order as the activities during each movie. We then administered a recognition test (see the SI Appendix, Section 3). Finally, all participants completed a vocabulary test (42), after which older adults completed the MMSE (41). The three-day retention interval was chosen based on pilot testing, to avoid floor and ceiling effects on the memory measures. For more methodological detail, see the SI Appendix (Sections 1.2 and 1.3).

### fMRI Data Analyses

Because we had strong *a priori* hypotheses regarding the brain regions that would be relevant in the RSA, we employed a ROI-based analytic strategy. Specifically, we selected the PMC and MTL parcels of the DN subsystems from the 17 networks/300 parcels cortex parcellation map of (28) to which we added ROIs of the left and right hippocampus. Following spatial preprocessing and prior to the RSA, data were normalized and detrended using second-order polynomials, spatially smoothed with a Gaussian kernel of 3-mm full-width at half maximum and Z-scored. To summarize the activity within each voxel during the period of interests in each run, we performed temporal compression by averaging the ninth to fourteenth scans (11.97 to 18.62 s) after the beginning of each activity. This temporal compression procedure resulted in one brain image for each activity, run, and participant (See SI Appendix Section 1.5 for additional details on image preprocessing for the RSA and Section 1.6 for a description of the mass univariate analyses). We then compared the similarity of the brain activity patterns in each parcel between the two runs. Finally, we computed reinstatement Z-scores that quantified the degree to which reinstatement activation patterns are more similar to their *matching* Day 1 activity encoding activation pattern than to the others (18, 19; see Figure 3 for details).

## Acknowledgements

We thank Ryan Kahle, Madeleine Schroedel, and Priscilla Mei for assistance with data collection and coding. We also thank Aaron B. Tanenbaum for his help with spatial preprocessing and selection of the fMRI sequences. This project was funded by NIH grant R21AG05231401 and supported in part by the Neuroimaging Informatics and Analysis Center (1P30NS098577). David Stawarczyk is currently supported by the European Union’s Horizon 2020 Research and Innovation Programme under the Marie SkłodowskaLCurie grant agreement No 798109. Joset A. Etzel was partially supported by the National Institutes of Health, grant number R37MH066078.

## Author contribution

DS, CW, JE, and JZ designed the study. DS performed the study. DS, JE, and JZ analyzed the data. DS, CW, and JZ wrote the paper with all other co-authors providing critical inputs. AS helped with the selection of the fMRI sequence and provided the spatial preprocessing pipeline used in the present work. All authors approved the final version of the manuscript.

## Supporting Information

### 1. Material and Methods

#### a. Participants

Participants were recruited from the Washington University School of Medicine Research Participant Registry, flyers posted on campus, and word of mouth. Potential participants first completed a prescreening interview over the phone. In addition to MRI contraindications, we excluded anyone who reported a history of neurological or psychiatric disorders, taking medication that could affect their cognitive functioning, or not having normal or corrected to normal vision and audition. The two age groups did not significantly differ in years of education [*t*(60) = −1.51, *p* = .14, *d* = .38; *M* = 15.85, *SD* = 1.71, for the young adults; *M* = 16.64, *SD* = 2.39, for the older adults] or percentage correct on the Shipley Institute of Living Scale (1) vocabulary test [*t*(60) = −1.88, *p* = .07, *d* = .48; *M* = 83.90%, *SD* = 8.33, for the young adults; *M* = 87.88%, *SD* = 8.28, for the older adults]. Each participant received $25.00 per hour for participating in the study. Nine other participants took part in the study but were excluded from the analyses for the following reasons: two young adults interrupted the study during scanning, technical issues lead to unusable data for two older adults, and five more older adults did not comply with the task instructions.

#### b. Materials

Each of the 45 activities comprised two parts: an initial *cue* segment that was identical for the A and B versions and lasted 6 s, followed by a *post-divergence* segment that sometimes included the changed feature and lasted 12 s with the last second including a fade to black transition. For some of the activities, the changed feature was an object that the actor contacted (e.g., pouring a glass of *milk* or a glass of *water*). For other activities, the changed feature was the action itself (e.g., doing *leg stretches* or *sit-ups* on a yoga mat). In all cases the changed feature was central to the activity performed. The critical manipulation was whether the post-divergence segment was the same in both movies (repeated activities), or whether that segment changed from the Day 1 to Day 2 movie (changed activities; e.g., the A version in Day 1 and the B version in Day 2). We included twice as many changed as repeated activities (i.e., 30 vs. 15) to increase the power to detect differences in neural activity associated with change processing.

Colored square-wave gratings were overlaid on the movies during the post-divergence segment. The gratings had a spatial frequency of approximately half a cycle per degree of visual angle and an opacity of 60%. The gratings gradually appeared over the movies during the first 1.5 s of the post-divergence segment (see **Figure S7**). Half of the gratings were red and vertical and half were green and horizontal. Gratings were presented in a fixed pseudo-random sequence with the constraint that they did not repeat for more than four consecutive activities. For clips that included a changed feature in the Day 2 movie, the gratings also changed; for clips that included a repeated feature in the Day 2 movie, the gratings remained the same^1^.

To counterbalance the assignment of activities to conditions, we created 12 experimental formats. We divided the 45 activities into 3 groups of 15 and rotated the groups through conditions across participants, such that each participant viewed two groups of changed activities and one group of repeated activities. The sequence of activities was fixed, beginning with the actor waking up and ending with her going to bed. The assignment of activities to experimental conditions (which ending was presented on Day 1, and whether the item was repeated or changed) was fixed in a pseudo-random sequence such that each third of the task (15 activities) contained five repeated activities with no more than six consecutive changed activities throughout the task. We also alternated the gratings and whether the A or B version of each activity appeared in the Day 1 movie. All stimulus materials were presented using E-Prime 2 software (2).

#### c. Procedure and task description

Participants completed the task in two sessions separated by three days (**Figure 1**). During the first session, participants watched both movies while lying in the scanner. Before the first functional run, we informed participants that they would watch a movie of an actor performing a series of everyday activities throughout the course of her day. We instructed participants to pay attention to her actions and the objects that she contacted. We called this functional run the “Day 1” viewing. To encourage attention during encoding of post-divergence segments, we instructed participants to associate the movie content with the colored gratings that we told them would also appear briefly after the onset of each activity. In the Day 1 movie, the initial cue segment of each activity was followed immediately and seamlessly by the post-divergence segment. Following each post-divergence segment, a fixation cross appeared in the middle of the screen for a mean duration of 4630 ms (*SD* = 10, range = 4585-4639 ms), with the onset of the cue segment for the next activity being synchronized with the onset of the next scan. This fixation cross provided a demarcation between events, which was necessary for the subsequent reinstatement task. It also allowed the blood-oxygen-level-dependent (BOLD) response to decrease before the onset of the next activity and permitted us to use the same time window across activities regarding the onsets of the video clips in the representational similarity analyses (RSA; see the fMRI analyses section for details). We presented two example activities in the scanner before the beginning of the first run to familiarize participants with the task.

After participants viewed the Day 1 movie, field map images and a high-resolution anatomical image were collected, taking approximately 10 minutes. We then told participants that their next task would be to watch another movie that depicted the same actor performing activities on another fictive day that occurred one week later in her life. We called this second functional run the “Day 2” viewing (see **Figure 1**). We explained to participants that the activities would appear in the same order as in the first movie, but that the clips would stop after a few seconds so that they could mentally replay the upcoming action viewed during the Day 1 movie. Participants were told that during this “reinstatement phase” they should imagine the event ending from the Day 1 movie in as much detail as possible. During the Day 2 viewing, each clip stopped after the cue segment, and a question appeared for 12 s (to match the duration of the Day 1 post-divergence segment) asking participants what happened next during Day 1. After this 12 s reinstatement phase, participants watched the post-divergence segment, during which participants were asked to pay specific attention to whether the activity features repeated or changed from Day 1. After each post-divergence segment, a 2 s fixation cross appeared, followed by two questions. The first question asked whether participants thought that they had correctly remembered the Day 1 post-divergence segment during the reinstatement phase. The second question asked whether the action in the post-divergence segment was repeated or changed. Participants used a button box to respond “yes” or “no” to the first question and “repeated” or “changed” to the second. The duration of each question was self-paced with a cut-off of 5 s, and a 250 ms fixation cross appeared between questions. As for the Day 1 movie, a fixation cross appeared at the end of the trial for a mean duration of 4668 ms (*SD* = 394, range = 4012-5345 ms) with the onset of the following cue segment being synchronized with the onset of the next scan.

After the second functional run we collected a second set of field map images and a high-resolution T2-weighted image. We then informed participants that Session 1 was over. We did not mention that Session 2 would consist of a memory test, in order to reduce the likelihood that participants would rehearse the activities during the three-day delay.

During Session 2, outside of the scanner, we first tested participant memory for the previously viewed activities using a cued recall task (see **Figure 1**). During the task, the questions appeared in the same order as the activities during each movie. For each activity, we first asked participants to recall features that appeared in the Day 2 clips by typing their response. For example, for the question, “What did the actor eat for breakfast?”, participants might respond “A banana.” The tested features were all critical features that varied between the A and B versions of the activities. After each response, we asked participants whether the way that the actor performed the activity changed from Day 1 to Day 2. Participants pressed the “1” key to indicate that the activity repeated exactly and the “2” key to indicate that the activity changed. When participants indicated that the activity had changed, they were prompted to type in the original Day 1 feature. For each of these questions, we asked participants to rate the confidence in their answer on a Likert scale ranging from 1 “Low” to 5 “High” (these confidence ratings are not relevant to the hypotheses investigated here, so we do not discuss them further). All responding during the task was self-paced. Following the cued recall task, we administered a recognition test of Session 1 activities (see **Section 3** of the Supplementary Information for details). After the recognition test, all participants completed a vocabulary test (1), and older adults completed the MMSE (3) last.

#### d. fMRI data acquisition

All fMRI data were collected using a 3 Tesla scanner (Magnetom Prisma, Siemens Erlangen, Germany) equipped with a 64-channel receiver head coil. Head movement was minimized with foam padding. Anatomical imaging including a multi-echo, T1-weighted MP-RAGE scan (TR = 2500 ms, TE = 1.81, 3.60, 5.39, and 7.18 ms, FOV 256 × 240 mm, matrix 320 × 300 × 208, voxel size 0.8 × 0.8 × 0.8 mm) and a high-resolution T2-weighted SPACE scan (TR = 3200 ms, TE = 564 ms, FOV 256 × 240 mm, matrix 320 × 300 × 208, voxel size 0.8 × 0.8 × 0.8 mm). BOLD fMRI was acquired using a T2*-weighted, multi-band accelerated EPI pulse sequence developed at the Center for Magnetic Resonance Research (CMRR) at the University of Minnesota (MB factor = 4, TR = 1330 ms, TE = 38.8 ms, FA 63°, matrix size 110 × 110 × 60, voxel size 2.4 × 2.4 × 2.4 mm, A >> P phase encoding). Whole brain coverage was obtained with 60 2.4 mm slices without in-plane acceleration (iPAT = 0). A mean of 782 (*SD* = 3.83, range: 775-800) and 1343.50 (*SD* = 29.78, range: 1285-1420) volumes were acquired in the first and second functional runs, respectively. Spin echo field maps were acquired after each fMRI run. Stimuli were displayed on a screen positioned at the rear of the scanner that participants viewed via a mirror mounted on the head coil.

#### e. Preprocessing of fMRI data

The functional data were analyzed in (2 mm isotropic) 711-2B atlas space (4). Atlas transformation was initially computed by composition of affine transforms (fMRI functional volume mean → T2w → T1w → 711-2B space representative target image). The last transformation step was refined by non-linear registration of each individual’s T1w to the atlas representative target using the FNIRT module in fsl (5, 6). One step final resampling of the functional data in atlas space combined retrospective head motion correction, magnetization inhomogeneity distortion correction via topup (7) and non-linear atlas transformation.

Because we had strong *a priori* hypotheses regarding the brain regions that would be relevant in the RSA, we employed a ROI-based analytic strategy. Specifically, we selected the PMC and MTL parcels of the DN subsystems from the 17 networks/300 parcels cortex parcellation map of Schaefer et al. (8). As this parcellation does not map subcortical areas, we added ROIs for two parcels for the left and right hippocampus, extracted using the Oro.nifti package (9) in R software (10). These hippocampal parcels were obtained from the minimal preprocessing pipelines for the Human Connectome Project’s (11) cifti subcortical structure using the cifti subcortical structure using the “-cifti-separate” Connectome Workbench function (12). We resampled the parcellation to atlas space using the 3dresample AFNI command (13) and warped it from MNI to our template using the “Old Normalise” command in Statistical Parametric Mapping (SPM 12 with updates *6685*).

Following spatial preprocessing and prior to the RSA, data were detrended using second-order polynomials with the 3dDetrend AFNI command, spatially smoothed (14) with a Gaussian kernel of 3-mm full-width at half maximum in SPM, and z-scored with PyMVPA (15). To summarize the activity within each voxel during the period of interests in each run, we performed temporal compression by averaging the ninth to fourteenth scans (11.97 to 18.62 s) after the beginning of each activity using the 3dTstat AFNI command. This averaging window corresponded to the post-divergence segments in the first run and reinstatement phases in the second run. It was chosen to account for the hemodynamic delay, and to reduce the chance of contamination from the blood-oxygen-level-dependent (BOLD) response associated with the fixation cross of the first run and post-divergence segment of the second run, since these started 18 s after the onset of the cue segment. This temporal compression procedure resulted in one brain image for each activity, run, and participant.

We then compared the similarity of the brain activity patterns in each parcel between the two runs. Specifically, a similarity matrix was constructed for each parcel within each participant by placing the 45 activities from the first run along one axis and the same 45 activities from the second run along the other axis, filling the matrix with all possible pairwise Pearson correlations (i.e., each Day 1 post-divergence segment correlated with each Day 2 reinstatement). If participants successfully reinstated the pattern of neural activity experienced while watching the post-divergence segments of the first run during the reinstatement phases of the second run, then we would expect the correlations along the diagonal (i.e., watching and reinstating the same activity) to be higher than off-diagonal correlations (i.e., watching and reinstating different activities). This tendency was quantified for each activity as its on-diagonal cell minus the average of the off-diagonal cells (after Fisher’s r to z transformation; 31, 56). To ensure the scores were comparable at the group level and not biased by possible off-diagonal matrix structure, each difference score was transformed by means of a dataset-wise permutation test (18) within each individual to form the final reinstatement Z-score used in all further analyses. Specifically, 1,000 unique random permutations of the Day 2 activity order were generated, and the difference score calculated for each of these permutations in each matrix, creating a separate null distribution for each participant and parcel. The reinstatement Z-score was the ranking of the actual (unpermuted) difference score in its null distribution, transformed to a Z-score. This was done using the R command qnorm(1/1001) resulting in a Z-score of 3.09 when the unpermuted score was larger than that from all permutations.

#### f. Mass univariate fMRI modeling

Following preprocessing, brain images were spatially smoothed with a Gaussian kernel of 6-mm full-width at half maximum in SPM. For each participant, BOLD responses during the Day 2 run were modeled at each voxel, using a general linear model (GLM). The post-divergence segments of repeated and changed items were modeled separately as epoch-related responses (lasting 12 s) and convolved with the canonical hemodynamic response function to create the regressors of interest. The cue segments and reinstatement phases were also modeled as epoch-related responses (lasting 6 s and 12 s, respectively), each with a single regressor across all trials. The two questions ending each trial were also modeled as epoch-related responses with a single regressor across all conditions. The design matrix included the realignment parameters to account for any residual movement-related effect, and a high-pass filter using a cutoff period of 128 s to remove low frequency drifts from the time series. Serial autocorrelations were estimated with a restricted maximum likelihood algorithm with an autoregressive model of order 1 (+ white noise). Based on this design, we computed a linear contrast to examine the brain regions more active in changed than repeated post-divergence segments. The contrasts of interest were first computed for each participant and were then entered in a random-effect one-sample *t*-test to assess the overall effect of change across all participants. Finally, we entered the contrasts in a two-sample *t*-test to assess group difference between young and older adults. For both contrasts, we report activations that were statistically significant using a threshold of *p* < .05, corrected for multiple comparisons (familywise error; FWE) over the whole brain or within a mask of the bilateral hippocampus.

To assess the possibility that brain activity while viewing the changed post-divergence segments could be modulated by event-specific reinstatement Z-scores, we computed the same model as described in the previous paragraph, but added a parametric regressor consisting of the trial-wise reinstatement Z-scores computed in the RSA, centered within each participant. Three models were computed, with either the mean PMC, MTL, or hippocampus reinstatement Z-scores as parametric regressor. We next computed the contrast of this parametric regressor against a baseline for each participant, which was entered in a random-effect one-sample *t*-tests to assess its overall effect across all participants. We then entered the contrasts in a two-sample *t*-test to assess the group difference between young and older adults. For both contrasts, we report activations that were statistically significant using a threshold of *p* < .05, corrected for multiple comparisons (FWE) over the whole brain or within a mask of the bilateral hippocampus.

Finally, to assess the neural response to the movies during Day 1 viewing, we also modeled BOLD responses at each voxel for each participant using a GLM. The 18 s of each activity were modeled separately as epoch-related responses and convolved with the canonical hemodynamic response function to create the regressor of interest. The design matrix also included the realignment parameters to account for any residual movement-related effect, and a high-pass filter was implemented using a cutoff period of 128 s to remove the low frequency drifts from the time series. Serial autocorrelations were estimated with a restricted maximum likelihood algorithm with an autoregressive model of order 1 (+ white noise). Based on this design, we computed a linear contrast to examine which brain regions were more active while viewing the movies than baseline. The contrasts of interest were first computed for each participant and were then entered in a random-effect one-sample *t*-tests to assess the overall effect movie viewing across all participants. We then entered the contrasts in a two-sample *t*-test to assess the difference between young and older adults. For both contrasts, we report activations that were statistically significant using a threshold of *p* < .05, corrected for multiple comparisons (FWE) over the whole brain. Only clusters with a size of *k* > 20 voxels are reported for each of these analyses.

#### g. Classifier MVPA

To further assess the effect of age on the neural response to the movies during Day 1 viewing, we computed the same GLM as in the previous sections using SPM, except that images were spatially smoothed with a Gaussian kernel of 3 (rather than 6)-mm full-width at half maximum. We then used the Pronto Toolbox (19) to determine whether a linear support vector machine classifier (*C* = 1) trained on the parameter estimate images corresponding to the movie regressor of every participant except one could accurately determine the age group of the remaining participant. We ran this analysis twice, using either a mask of the PMC or MTL. To assess whether the classification accuracy between the two groups was above chance, a permutation procedure randomly shuffling the group label of the parameter estimate images was used (1000 permutations using the default Pronto settings). Given the unequal number of participants in each age group, we report the balanced accuracy values that give equal weight to the accuracies obtained for each group.

#### h. Behavioral performance modeling

All analyses of memory performance and reinstatement effects were computed in R software (10) using linear or logistic mixed effect models with subjects and activities as random effects. Logistic models were used when the dependent variable was binary. Models were fitted using functions from the lme4 package (20), hypothesis tests were performed using the Anova function of the car package (21), and post hoc comparisons using the Tukey method were conducted using the emmeans function from the emmeans package (22). Comparisons between nested models were performed with the anova function of the lme4 package. Centering of the reinstatement Z-scores was done using the center function of the misty package (23). Finally, the plot_model and get_model_data functions from the sjPlot package (24) were used to compute parameter estimates (i.e., model estimated probabilities) for description and visualization.

##### i. Cued recall response coding

All open-ended responses made during the cued recall task (i.e., answers to the questions about Day 2 activity features and about Day 1 activity features for activities classified as changed) were coded by the first author and a second independent rater. Participants’ Day 2 recall responses were classified as one of three types. *Correct Day 2 recall* responses were correct descriptions of the criterial Day 2 activity feature. *Day 1 intrusions* were responses that included the criterial Day 1 feature that did not appear on Day 2. All remaining responses were considered as *Incorrect*. The Cohen’s κ (25) between raters was .85. When participants classified activities as changed on the Day 2 recall test, they were asked to recall the Day 1 feature. The same raters coded these responses as *correct Day 1 recalls* or *incorrect Day 1 recalls* depending on whether the participants recalled the critical Day 1 activity feature or not, respectively. Cohen’s κ for Day 1 recalls was .83.

### 2. Results and Discussion

#### a. Behavioral performance

##### i. Self-reported reinstatement success and change detection during Day 2 viewing

During Day 2 viewing, each activity was paused after the cue segment, and participants were asked to mentally replay how the activity ended on Day 1. After seeing the changed or repeated post-divergence segment, they reported whether their reinstatement was successful and whether the Day 2 ending was repeated exactly or included a changed feature (see **Figure 1**). To characterize subjective accuracy in reinstating Day 1 features and objective accuracy in activity type classifications (see **Table S3**), we fitted 2 (Activity Type: Repeated vs. Changed) × 2 (Age Group: Young vs. Older) models to each measure. The model for reinstatement success indicated that older adults reported significantly more successful Day 1 reinstatements than young adults [χ*^2^*(1) = 4.55, *p* = .03]. In contrast, the model for classification accuracy showed the opposite pattern: young adults showed significantly greater classification accuracy than older adults [χ*^2^*(1) = 6.68, *p* = .01]. No other effects were significant [*largest* χ*^2^*(1) = 1.10, *p =*.30]. Together, these results suggest that metacognitive accuracy about memory for Day 1 features was better for young than older adults. Relevant to our interest in age differences in change processing, these findings imply that reinstatement success should be less predictive of change detection accuracy for older than young adults.

To determine the extent to which successful reinstatement of Day 1 activity features contributed to detection of changed Day 2 features for both age groups, we examined the relationship between reinstatement success and classification accuracy for changed activities only. Finding that successful reinstatement predicts more accurate change detection would be consistent with the proposal that prior event retrieval facilitates detection of changed event features. Further, finding that such an effect is greater for young than older adults would suggest that older adults are impaired in their ability to use of prior event retrievals to update their memories. A 2 (Reinstatement: Successful vs. Unsuccessful) × 2 (Age Group: Young vs. Older) model (see **Table S4**) indicated a significant effect of Reinstatement [χ*^2^*(1) = 25.95, *p* < .001] and a significant Reinstatement × Age Group interaction [χ*^2^*(1) = 9.97, *p* = .002], showing that successful reinstatements were associated with more accurate change detection for young adults (*z* ratio = 5.94, *p* < .001) but not for older adults (*z* ratio = 1.00, *p* = .32). These results suggest that prior event retrievals enabled more effective change detection for young than older adults.

Next, we assessed the extent to which reinstatement success and change detection during Day 2 viewing were associated with subsequent memory performance in the cued recall task. Prior behavioral work has shown that accurate change detection is associated with better cued recall for Day 2 changed activity features and predicts overall rates of change recollection for both young and older adults (26). Here, we examined whether those findings would replicate, and we extended this analysis to the self-report measure of reinstatement. Given the age differences in the relationship between reinstatement success and change detection above, we expected age differences in the associations between these measures and behavioral memory performance.

**Figure S8** displays Day 2 recall and change recollection measured on the cued recall test conditionalized on reinstatement success (left panels) and change detection accuracy (right panels) measured during Day 2 viewing. To examine how the Day 2 measures predicted performance on behavioral memory measures for changed activities at test, we fitted separate models to each test measure including the Day 2 measure (reinstatement or change detection) and age group as fixed effects. The model testing the association between reinstatement success and Day 2 recall (top left panel) indicated that successful reinstatement was associated with more accurate Day 2 recall [χ^2^ (1) = 12.49, *p* < .001] and this effect did not interact with Age Group [χ^2^ (1) = 0.14, *p* = .71]. Reinstatement was also associated with greater change recollection for both age groups (middle left panel) [χ^2^ (1) = 61.93, *p* < .001] and this effect did not interact with Age Group [χ^2^ (1) = 0.05, *p* = .82]. The same models including change detection accuracy (instead of reinstatement) as the fixed effect indicated no association between change detection and Day 2 recall (top right panel) [χ^2^ (1) = 3.44, *p* = .07], and a positive association between change detection and change recollection (middle right panel) [χ^2^ (1) = 87.85, *p* < .001]. These effects did not interact with Age Group [*largest* χ^2^ (1) = 0.74, *p* = .39]. The lack of association between change detection and Day 2 recall accuracy fails to replicate earlier findings (26), but the positive association between reinstatement and Day 2 recall implicates a role for prior event retrieval in the updating of memory to include changed features. Also, the strong positive associations with change recollection for both reinstatement and change detection are not surprising given that all these measures are considered to assay the accessibility of Day 1 activity features during various phases of the experiment.

The findings that self-reported reinstatement and change recollection were both positively associated with Day 2 recall and each other suggest that the association between successful Day 1 reinstatement and change recollection is a consequence of reinstatement enabling change detection and the formation of configural memory representations during Day 2 viewing. We examined this possibility by testing whether the association between reinstatement and Day 2 recall remained when change recollection was also included as a fixed effect. The rationale here is that if change recollection predicts Day 2 recall when controlling for reinstatement, but reinstatement does not predict Day 2 recall when controlling for change recollection, then this would imply a mediating role for reinstatement during Day 2 viewing in the positive association between change recollection and Day 2 recall at test. **Figure S9** displays Day 2 recall accuracy predicted by reinstatement success and change recollection controlling for the effects of the other variable. A 2 (Reinstatement: Successful vs. Unsuccessful) × 2 (Change Recollection: Recollected vs. Not Recollected) × 2 (Age Group: Young vs. Older) model indicated no significant effect of Reinstatement [χ*^2^*(1) = 0.49, *p* = .48] and significant effects of Change Recollection [χ*^2^*(1) = 219.95, *p* < .001], and Age Group [χ*^2^*(1) = 7.18, *p* = .008]. There were no significant interactions [*largest* χ*^2^*(1) = 3.76, *p* = .052].

Together, these results indicate that self-reported reinstatement of Day 1 features during Day 2 viewing was associated with better cued recall of the changed Day 2 features in both age groups. However, this effect did not remain significant after controlling for the ability to recollect change. This suggests that the association between reinstatement of Day 1 experiences during viewing of Day 2 activities and recall of those Day 2 activities is mediated through recollecting at test that the activity had earlier changed. However, the current design does not allow for strong causal conclusions about this potential mediation.

##### ii. Day 1 intrusions during Day 2 recall

Analyses of Day 1 intrusions during Day 2 recall (**Figure S10**) revealed results that mostly mirrored those of correct recall. We fitted a model to Day 1 intrusions that was comparable to the model for Day 2 recall. It included a fixed effect of Activity Type that included one level for all repeated activities and three levels of changed activities that corresponded with each change classification type, as well as a fixed effect of Age Group. The model indicated a significant effect of Activity Type [χ*^2^*(3) = 144.00, *p* < .001], no significant effect of Age Group [χ*^2^*(3) = 0.12, *p* = .73], and no significant Activity Type × Age Group interaction [χ*^2^*(3) = 2.26, *p* = .52]. For both age groups, recollecting change was associated with fewer intrusions compared to baseline rates for repeated items (*z* ratio = −3.10, *p* = .01). This is not surprising given that this conditional cell for changed activities comprises instances when participants reported the Day 1 feature twice, presumably because they were guessing. Intrusion rates for changed activities did not differ between when changes were remembered but the Day 1 feature was not recalled and when changes were not remembered (*z* ratio = −0.92, *p* = .79). Both conditional cells were associated with intrusion rates above baseline (*smallest z* ratio = 7.56, *p* < .001).

We also examined whether Day 1 intrusions during Day 2 recall of changed activities were predicted by reinstatement success and change detection during Day 2 viewing (see **Figure S8,** bottom panels). The model with Age Group and Reinstatement as fixed effects (left panel) revealed no significant effects of Reinstatement [χ*^2^*(1) = 0.81, *p* = .37], or Age Group [χ*^2^*(1) = 1.31, *p* = .25], but the Reinstatement × Age Group interaction was significant [χ*^2^*(1) = 4.80, *p* = .03]. Successful reinstatement was associated with fewer intrusions for young adults (*z* ratio = 2.01, *p* = .04) but not for older adults (*z* ratio = −1.23, *p* = .22). The comparable model including Change Detection as a fixed effect (right panel) indicated no significant effects of Change Detection [χ*^2^*(1) = 0.02, *p* = .90], or Age Group [χ*^2^*(1) = 1.03, *p* = .31], and no significant Change Detection × Age Group interaction [χ*^2^*(1) = 0.01, *p* = .91].

It may seem somewhat surprising that older adults did not produce more Day 1 intrusions than young adults. Several theories of age-related episodic memory deficits assume that older adults’ impaired memory stems from a higher susceptibility to interference (e.g., 27). However, there is mixed empirical evidence supporting this claim (for a review, see 28). One possibility is that the lack of an age-related difference in the present study reflects younger adults having access to more activity features and a more liberal reporting criterion than older adults (for similar arguments, see 29). Another possibility is that the shorter delay between Day 2 and test used here (72 hours) compared to that used in previous behavioral work (one week; 15) did not allow enough time for older adults’ deficit in source monitoring of intrusions to emerge.

To further investigate these results, we also examined the specific kinds of errors that the participants from each group made during the Day 2 recall of changed activities that were not Day 1 intrusions. To do so, we classified the remaining errors as *No Response* (i.e., “I don’t remember” and similar answers), *Ambiguous* (i.e., the participants gave an answer that was not specific enough to discriminate between the two versions of the activity), *Inter-Activity Intrusion* (i.e., participants recalled a feature from a different activity), and *Other* (i.e., participants recalled an extra-experimental activity feature). As an illustration of these response types, consider when the actor opened the refrigerator to get water on Day 1 and milk on Day 2. An *Ambiguous* response could be “cold drink,” an *Inter-Activity Intrusion* could be “Gatorade” if that was the criterial feature of another activity, and *Other* could be “Wine” if that was not the criterial feature of an activity. For young adults, of the 218 incorrect Day 2 recalls of changed activities that were not Day 1 intrusions, 25% were *No Response*, 21% were *Ambiguous*, 14% were *Inter-Activity Intrusions*, and 40% were *Other*. For older adults, of the 244 incorrect Day 2 recalls to changed activities that were not Day 1 intrusions, 28% were *No Response*, 14% were *Ambiguous*, 16% were *Inter-Activity Intrusions*, and 43% were *Other*. Thus, it seems that the distributions of error types (other than Day 1 intrusions) were comparable for young and older adults.

#### b. RSA analyses

##### i. Change detection accuracy and Day 1 intrusions: Between-participant associations with mean PMC and MTL reinstatement Z-scores

To examine between-participant differences in the relationship between neural reinstatement during Day 2 viewing and change detection in the scanner, we computed mean between-participant correlations between RSA reinstatement Z-scores and *Change detection accuracy* (see **Figure S11,** middle panels). These analyses revealed no significant effect of reinstatement Z-scores in the either the PMC or MTL [*largest* χ*^2^*(1) = 2.18, *p* = .14], but the interactions with Age Group were significant [χ*^2^*(1) = 4.69, *p* = .03, for the PMC; χ*^2^*(1) = 8.53, *p* = .003, for the MTL]. Higher mean reinstatement Z-scores in both regions predicted more accurate change detection for young adults [χ*^2^*(1) = 3.99, *p* = .046, for the PMC; χ*^2^*(1) = 10.60, *p* = .001, for the MTL] but not for older adults [χ*^2^*(1) = 0.75, *p* = .39, for the PMC; χ*^2^*(1) = 0.15, *p* = .70, for the MTL]. These findings are consistent with the behavioral results showing that self-reported reinstatement success predicted better change detection for young but not older adults. The comparable model predicting *Day 1 intrusions* (see **Figure S11**, right panels) indicated no significant effects of reinstatement Z-scores in either the PMC or MTL and no significant interaction with Age Group [*largest* χ*^2^*(1) = 1.92, *p* = .17].

##### ii. Between-participant analyses of individual parcel reinstatement Z-scores

Analyses of the relations between the mean PMC and MTL reinstatement Z-scores revealed several significant associations (see main text). However, it is unclear whether these associations were driven by specific parcels above and beyond the effects of the other parcels in each of the two areas and whether some significant associations might have been obscured by different parcels within the PMC or MTL having opposite associations with memory performance. To investigate these possibilities, we used logistic mixed effect models with between- and within-participant reinstatement Z-scores, as in the main manuscript, but added the individual Z-scores of each parcel as fixed effects rather than a single mean value representing the average of all the parcels. We then used the anova function of the lme4 package (20) to determine whether adding these single parcel Z-scores relative to a baseline model comprising only the random effects improved model fits. We then examined whether these model fits were further improved by adding the interaction terms that included Age Group as a fixed effect.

First, regarding *Day 2 recall accuracy* for changed activities, adding the between-participant reinstatement Z-scores for the PMC parcels did not improve model fit, nor did adding the interaction terms with Age Group [*largest* χ (12) = 16.31, *p =* .18]. These results indicate that the significant effect of the mean PMC reinstatement Z-score reported in the main manuscript was not driven by specific parcels. We found similar results for the MTL: the model fit was not improved by adding the reinstatement Z-scores [χ (6) = 11.76, *p* = .07], or the interactions with Age Group [χ (6) = 5.68, *p* = .46]. We next performed the same analyses for *Day 1 intrusions* for changed activities. The model fit was not improved by adding reinstatement Z-scores for the PMC or MTL parcels or the interactions with Age Group [*largest* χ (12) = 12.80, *p =*.38].

Second, regarding *change recollection* for changed activities in the cued recall task, adding the between-participant reinstatement Z-scores for the PMC parcels did not improve model fit [χ (12) = 19.64, *p* = .07], but adding the interaction terms with Age Group did [χ (12) = 23.29, *p* = .03]. Further analyses showed that individual parcel reinstatement Z-scores improved model fit for the young [χ (12) = 25.69, *p* = .01] but not the older adults [χ (12) = 15.28, *p* = .23]. For the young adults, higher reinstatement Z-scores were associated with higher change recollection accuracy in parcels 115 [χ^2^ (1) = 20.25, *p* < .001] and 292 [χ^2^ (1) = 5.18, *p* = .02]. No significant effect was found in the models including the MTL parcels [*largest* χ (6) = 11.80, *p =* .07].

Third, for *self-reported reinstatement success* during Day 2 viewing, adding the between-participant reinstatement Z-scores for the PMC parcels did not significantly improve model fit [χ*^2^*(12) = 17.39, *p* = .14], but adding the interaction terms with Age Group did [χ*^2^*(12) = 24.14, *p* = .02]. Further analyses showed that the individual parcel Z-scores improved model fit for the young [χ*^2^*(12) = 25.13, *p* = .01] but not the older adults [χ*^2^*(12) = 18.16, *p* = .11]. For the young adults, higher reinstatement Z-scores were associated with higher self-reported reinstatement accuracy in parcel 115 [χ*^2^*(1) = 17.31, *p* < .001], whereas higher reinstatement Z-scores were associated with lower self-reported reinstatement accuracy in parcel 114 [χ*^2^*(1) = 4.32, *p* = .04]. No significant effects were found in the models including the MTL parcels [*largest* χ*^2^*(6) = 8.15, *p =* .23].

Finally, regarding *change detection accuracy* during Day 2 viewing, model fits were improved by adding the between-participant reinstatement Z-scores for the PMC parcels [χ (12) = 21.10, *p* = .049], and the interaction terms with Age Group [χ (12) = 30.41, *p* = .002]. Further analyses revealed that adding the parcels improved model fit for the older [χ (12) = 26.67, *p* = .009] and young adults [χ (12) = 25.07, *p* = .01]. However, reinstatement Z-scores in different parcels for each age group were associated with change detection performance: for the young adults, higher reinstatement Z-scores were associated with better change detection accuracy in parcels 115 [χ^2^ (1) = 14.39, *p* < .001] and 292 [χ^2^ (1) = 4.49, *p* = .03], whereas higher reinstatement Z-scores were associated with worse change detection accuracy in parcel 142 [χ^2^ (1) = 5.11, *p* = .02]. For the older adults, higher reinstatement Z-scores were associated with better change detection accuracy in parcel 142 [χ^2^ (1) = 18.72, *p* < .001] and with lower change detection accuracy in parcels 117 [χ^2^ (1) = 4.34, *p* = .04] and 275 [χ^2^ (1) = 11.73, *p* < .001]. In the models including the MTL parcels, adding the individual parcels did not improve model fit [χ (6) = 11.74, *p =* .07] but adding the interaction term with Age Group did [χ (6) = 12.65, *p* = .049]. Further analyses showed that adding the effect of the individual parcels improved model fit in the young [χ (6) = 14.45, *p* = .03] but not older adults [χ (6) = 6.83, *p* = .34]. In the young adults, higher reinstatement in parcel 144 was associated with better change detection [χ^2^ (1) = 4.26, *p* = .04].

Together, these results show that memory for the original Day 1 activity feature (measured by self-reported reinstatement success, change detection accuracy during Day 2 viewing, and change recollection accuracy on the cued recall task) was strongly associated with PMC reinstatement in parcel 115 (corresponding to the posterior cingulate cortex) in the young adult group. These results are concordant with past fMRI studies that examined memories for naturalistic stimuli using RSA and other multivariate analyses and showed that activity patterns in these regions are particularly predictive of subsequent memory in this population (for a review, see 17). This involvement of parcel 115 and both change detection and self-reported reinstatement success in only the young adults might explain the behavioral differences between the two age groups regarding the association between these two variables: older adults reported greater reinstatement success but, unlike the young adults, self-reported reinstatement did not predict better change detection during Day 2 viewing, possibly because change detection was associated with a specific set of parcels for older adults.

##### iii. Within-participant analyses of individual parcel reinstatement Z-scores

For *Day 2 recall accuracy* for changed activities in the cued recall task, adding the fixed effects of the PMC parcels improved model fit [χ*^2^*(12) = 25.27, *p* = .01] but adding the interaction term with Age Group did not [χ*^2^*(12) = 9.87, *p* = .63]. Examination of the parameter estimates for the individual parcels showed that higher reinstatement Z-scores were associated with higher Day 2 recall accuracy in parcels 291 [χ*^2^*(1) = 7.19, *p* = .007] and 292 [χ*^2^*(1) = 6.63, *p* = .01], whereas higher reinstatement Z-scores predicted lower Day 2 recall accuracy in parcels 141 [χ*^2^*(1) = 5.60, *p* = .02]. No significant effects were found for the models including the MTL parcels [*largest* χ*^2^*(6) = 10.11, *p =*.12]. For the remaining behavioral memory measures for changed activities, no significant effects were found for the models including either the PMC or MTL parcels for self-reported reinstatement success [*largest* χ*^2^*(12) = 16.08, *p =* .19], change detection accuracy [*largest* χ*^2^*(12) = 20.74, *p =* .054], change recollection accuracy [*largest* χ*^2^*(12) = 15.36, *p =* .22], or Day 1 intrusions [*largest* χ*^2^*(12) = 20.23, *p* = .06].

In sum, these analyses showed a more complex picture for within-than between-participant reinstatement Z-scores. As described in the main text, there were no positive associations between behavioral performance and mean within-participant reinstatement Z-scores in either the PMC or MTL. However, when examining the contribution of individual parcels above and beyond the others parcels in each area, within-participant strength of reinstatement Z-scores were associated with *better* memory for changed features for some PMC parcels (parcels 291 and 292 in the right retrosplenial cortex, Rsp), and was associated with *worse* memory for changed features in another PMC parcel (parcel 141 in the left Rsp). To the best of our knowledge, these results are the first to reveal that reinstatement in closely located areas within the same cortical region can have opposite effects on subsequent memory performance. Together, these results support the view that extended cortical areas such as the PMC comprise distinct subregions (8, 30, 31), and indicate that within-participant neural pattern reinstatement in some PMC parcels might be detrimental to memory for changed features, possibly by disrupting the creation of configural representations and thus leading to proactive interference (32). However, this is the first finding of this sort, so replication attempts are needed to draw firm conclusions.

##### iv. Conjoint associations of RSA reinstatement Z-scores and self-reported reinstatement success with Day 2 recall accuracy for changed activities

Because both mean neural reinstatement in the PMC and self-reported reinstatement success were positively associated with higher memory accuracy for the changed Day 2 activity features, and because neither effect remained significant after controlling for change recollection (see main text), we tested whether both measures of reinstatement were significantly associated with Day 2 recall of changed features above and beyond each other. These models included self-reported reinstatement success, mean PMC reinstatement Z-scores at the between-participant level, and Age Group as fixed effects (see **Figure S12** for Day 2 recall estimates by mean neural [left panel] and self-reported [right panel] reinstatement above and beyond the other effect). Both the PMC reinstatement Z-scores [χ*^2^*(1) = 4.27, *p* = .04], and self-reported reinstatement [χ*^2^*(1) = 10.99, *p* < .001] were significant in these models, but the interaction terms were not significant [*largest* χ*^2^*(1) = 2.83, *p =* .09].

These results showed that between-participant differences in PMC neural reinstatement Z-scores and self-reported reinstatement accuracy both independently predicted recall of changed Day 2 features. Although the effects of these two indices of reinstatement on memory for changed features were fully explained by change recollection, these results suggest that these two measures reflect partly distinct but complementary processes. A possibility is that participants mostly made their reinstatement judgments based on whether they were able to remember the specific features that changed between the two movies, whereas neural reinstatement in the PMC reflected the retrieval of more abstract features of the event models (e.g., spatio-temporal context within the movies of each activity, 33) but not necessarily the changed features. Another possibility is that seeing the activity endings led to feature activation that participants misattributed to reinstatement that had occurred before seeing the ending. This is a form of source confusion, and older adults are more susceptible to such misattributions (34). Future studies might illuminate this issue with more objective behavioral measures of reinstatement accuracy.

#### c. Hippocampal response for the difference between changed and repeated activities, reinstatement Z-scores, and behavioral performance

Models of memory updating propose that pattern completion based on previous memories leads to prediction errors when change is experienced, which can then drive new learning including integration processes to form configural memory representations (26, 35). To assess this possibility, we first examined whether participants who had higher mean reinstatement Z-scores in the MTL and PMC were also those who showed higher neural responses in the anterior hippocampal clusters that we found to be more activated while viewing changed as compared to repeated post-divergence segments (see **Figure S6**), and whether this association differed between the age groups. Then we examined whether participants with larger hippocampal responses had better behavioral performance on measures of memory for change.

To do so, for each participant we extracted the parameter estimates of the bilateral hippocampal clusters that were more activated during changed than repeated activity viewings from the random-effect one-sample *t*-test using Marsbar Toolbox (36). Given that these estimates in the left and right hippocampus were highly correlated (*r* = .79), we combined them into a single mean score for subsequent analyses. We then computed multiple regression models with hippocampal response as the dependent variable and reinstatement Z-scores in either the PMC or MTL, Age Group, and their interaction as predictor variables. As there was no difference in hippocampal activity between young and older adults, we do not report the effect of age here to avoid redundancy. For the MTL model, there was no effect of mean reinstatement Z-score [*F*(1, 58) = 0.005, *p* = .94, *η*^2^*_p_* < .001] and no significant interaction with Age Group [*F*(1, 58) = 0.04, *p* = .83, *η*^2^*_p_* < .001]. The PMC model indicated a similar lack of evidence for effects, as neither the effect of mean reinstatement Z-score [*F*(1, 58) < 0.01, *p* = .98, *η*^2^*_p_* < .001], nor the interaction with Age Group [*F*(1, 58) = 0.03, *p* = .87, *η*^2^*_p_* = .002], was significant.

We next examined whether the difference in hippocampal response for changed than repeated activities could predict behavioral performance. To do so, we used linear mixed effects models to examine whether adding the hippocampal response and its interaction with Age Group in the previously described models assessing the association between reinstatement Z-scores and behavioral performance improved model fits. Results showed that this was not the case in either region for Day 2 recall accuracy [χ*^2^*(2) = 2.41, *p* = .30, for the MTL; χ*^2^*(2) = 2.49, *p* = .29, for the PMC], Day 1 intrusions during Day 2 recall [χ*^2^*(2) = 1.10, *p* = .58, for the MTL; χ*^2^*(2) = 0.98, *p* = .61, for the PMC], change recollection [χ*^2^*(2) = 3.46, *p* = .18, for the MTL; χ*^2^*(2) = 3.62, *p* = .16, for the PMC], or self-reported reinstatement [χ*^2^*(2) = 2.82, *p* = .24, for the MTL; χ*^2^*(2) = 2.81, *p* = .25, for the PMC]. Unsurprisingly, given that hippocampal responses were not significantly related to reinstatement Z-scores, all the significant effects of reinstatement Z-scores described in previous sections and in the main text remained unchanged. However, participants with larger differences in bilateral hippocampal responses when viewing changed than repeated activity endings were more likely to correctly detect changes during Day 2 viewing. Adding the parameter estimates for the hippocampal response and their interaction with Age Group significantly improved fit for the MTL [χ*^2^*(2) = 9.95, *p* = .007] and PMC [χ*^2^*(2) = 9.07, *p* = .01] models. Results also showed that higher parameter estimates predicted better change detection [χ*^2^*(1) = 9.94, *p* = .002, for the MTL; χ*^2^*(1) = 8.64, *p* = .003, for the PMC] and that these effect did not interact with Age Group [*largest* χ*^2^*(12) = 1.07, *p =* .30]. The interaction between reinstatement Z-scores and Age Group described above at the between-participant level (see Section 2.2.1) remained significant [χ*^2^*(1) = 10.79, *p* = .001, for the MTL; χ*^2^*(1) = 5.80, *p* = .02, for the PMC]. Higher mean reinstatement Z-scores still predicted better change detection for young adults in the PMC [χ*^2^*(1) = 4.33, *p* = .04] and MTL [χ*^2^*(1) = 11.01, *p* < .001]. Finally, the within-participant interaction between change detection performance and mean PMC reinstatement Z-scores also remained significant [χ*^2^*(1) = 4.69, *p* = .03]. Changes in trials with higher reinstatement Z-scores were still less likely to be detected in the older adults [χ*^2^*(1) = 5.00, *p* = .03].

The association between the intensity of the difference in the hippocampal response to changed compared to repeated activities and change detection is concordant with previous studies showing that the anterior hippocampus is one brain region that is most consistently involved in novelty processing (for reviews, see 37, 38). However, this activity was not modulated by Age Group and was unrelated to neural reinstatement Z-scores and several behavioral measures (except for Day 2 recognition accuracy of changed activities, see section 3.5 below). Although these findings do not support our hypothesis that the hippocampal response would (at least partly) explain the association between reinstatement Z-scores and subsequent memory performances, they are nonetheless consistent with previous studies showing that anterior hippocampal engagement does not necessarily reflect successful encoding (e.g., 39). A possibility is that this anterior hippocampal activity solely reflects scene construction processes that are more strongly engaged during changed than repeated events because of the new perceived event features. The engagement of these scene construction processes would depend on change detection, but they would not facilitate subsequent episodic recollection, nor depend on how well participants were able to remember the original Day 1 event features during the reinstatement phases (37). Further studies should be conducted to further assess this possibility.

#### d. Classifier MVPA results

An unexpected finding of the RSA analyses was that reinstatement Z-scores in the PMC and MTL did not differ between age groups. To examine whether there might nonetheless have been age-related differences in how participants initially perceived the activities, we examined whether there were group differences in brain activity while encoding the activity features during Day 1 viewing. Across all participants, mass univariate analyses revealed the large set of brain regions typically activated while watching movie stimuli (see **Figure S13**). A two-sample *t*-test revealed that mass univariate activation levels did not differ between the two groups in any region (no cluster was activated above threshold). We next used a pattern classifier to examine whether the voxelwise distribution of activation in the PMC and MTL differed between young and older adults (see section 1.7 for a methodological description). The balanced accuracy for the classifier ran on activity patterns within the PMC mask was 74.79% (*p* = .001); 85.29% (*p* = .002) for young adults and 64.29% (*p* = .002) for older adults. The balanced classification accuracy for activity patterns within the MTL mask was 87.29% (*p* = .001); 85.29% (*p* = .002) for the young adults and 89.29% (*p* = .001) for the older adults. Thus, although young and older adults did not differ in the extent to which they were able to reinstate during the Day 2 reinstatement phases the neural activity patterns they had during the Day 1 viewing, these Day 1 neural activity patterns nonetheless differed between the two groups.

### 3. Recognition task

#### a. Task description

Following the cued recall task, we tested memory for Session 1 activities using a recognition task (see **Figure S14**). Activities were presented in the same order as in the cued recall test (and the Day 2 movie). We asked participants to choose which of two still shots appeared in the Day 2 movie. Each shot showed a key frame featuring the criterial feature from the A or B version of the activity without gratings. The pictures appeared on opposite sides of the screen. The assignment of activity version to the left or right side was constrained such that the same version did not appear on the same side of the screen more than three times consecutively. Participants chose the left shot by pressing the “1” key and the right shot by pressing the “2” button. The unchosen shot then appeared in the center of the screen, and participants were instructed to indicate whether the activity had appeared in the Day 1 movie. They responded “yes” by pressing the “1” button (to indicate that the activity had changed) and “no” by pressing the “2” button (to indicate that the activity had repeated). Participants rated their confidence in the accuracy of each response on the same Likert scale as in the cued recall task (ranging from 1 “Low” to 5 “High”). All responses were self-paced.

#### b. Behavioral results

To examine Day 2 recognition accuracy for the recognition task, we first fitted a model with Activity Type (Repeated vs. Changed) and Age Group (Young vs. Older) as fixed effects. Consistent with the cued recall task, **Figure S15** shows that Day 2 recognition accuracy was significantly higher for repeated than changed activities [χ*^2^*(1) = 16.29, *p* < .001] and that younger adults significantly outperformed older adults [χ*^2^*(1) = 9.03, *p* = .003]. There was no significant Activity Type × Age Group interaction [χ*^2^*(1) = 0.50, *p* = .48]. Next, we fitted a similar model to only Changed activities with those activities conditionalized on whether participants remembered change. Results revealed that changed activities for which participants reported that both versions had appeared in Session 1 (and thus remembered that a feature changed) were associated with significantly higher Day 2 recognition accuracy than changed activities for which participants did not indicate seeing both versions in Session 1 [χ*^2^*(1) = 61.95, *p* < .001], and this effect did not interact with Age Group [χ*^2^*(1) = 0.81, *p* = .37]. Finally, older adults reported remembered change significantly less often than young adults (Older = .64, *95% CI* = [.54, .73]; Younger = .76, *95% CI* = [.69, .82]), [χ*^2^*(1) = 5.82, *p* = .02]). Thus, as in the cued recall task, both age groups showed similar positive associations between memory for Day 2 activities and remembering change, but older adults experienced this association for fewer activities.

We next examined whether self-reported reinstatement success and change detection predicted better recognition performance for changed activities. To do so, we fitted models to Day 2 recognition accuracy for changed activities that included Age Group along with either self-reported reinstatement success or change detection accuracy as fixed effects. The model including self-reported reinstatement success (**Figure S16,** top left panel) indicated no significant effect of Reinstatement [χ*^2^*(1) = 2.51, *p* = .11], and a significant effect of Age Group [χ*^2^*(1) = 7.46, *p* = .006], that were qualified by a significant interaction [χ*^2^*(1) = 6.44, *p* = .01], showing that self-reported reinstatement success was associated with higher Day 2 recognition accuracy for young (*z* ratio = 2.80, *p* = .005) but not older (*z* ratio = 0.10, *p* = .32) adults. **Figure S16** (bottom left panel) shows that self-reported reinstatement success was also associated with better memory that activities changed in Session 1 [χ*^2^*(1) = 12.76, *p* < .001]. This effect did not significantly interact with Age Group [χ*^2^*(1) = 0.23, *p* = .63]. The model including change detection (**Figure S16,** top right panel) indicated no significant effect of Change Detection [χ*^2^*(1) = 2.63, *p* = .10], a significant effect of Age Group [χ*^2^*(1) = 5.87, *p* = .02], and no significant Change Detection × Age Group interaction [χ*^2^*(1) = 1.31, *p =* .25]. However, **Figure S16** (bottom right panel) shows that accurate change detection was associated with better memory that activities changed in Session 1 [χ*^2^*(1) = 39.22, *p* < .001], there was a significant effect of Age Group [χ*^2^*(1) = 4.92, *p* = .03] showing better memory or change for young than older adults, and there was no significant Change Detection × Age Group interaction [χ*^2^*(1) = 0.44, *p* = .51].

Together, these results closely parallel those of the cued recall task, with the main difference being that, for older adults, self-reported reinstatement success did not predict correct recognition of changed Day 2 activities, whereas it did predict higher recall accuracy for the changed Day 2 features (most likely because of between the two tasks differences regarding how memory is assessed, see the last paragraph of the next section for a brief discussion of this point). As in the cued recall task, change detection accuracy was more strongly related to memory that a change occurred than to recognition of the changed Day 2 features for both age groups.

#### c. RSA results

We next tested our hypothesis that higher neural reinstatement Z-scores would predict better memory for changed activity features. As with the cued recall task, we first fitted models with the mean between- and within-participant reinstatement Z-scores in the PMC or MTL and Age Group as fixed effects. Recognition performance predicted by mean between- and within-participant neural reinstatement Z-scores appear in **Figure S17.** Analyses of *Day 2 recognition accuracy* for changed activities (left panels of **Figure S17**) revealed no significant effect of mean PMC or MTL reinstatement Z-scores, either between- or within-participants, and no significant interaction with Age Group [*largest* χ*^2^*(1) = 2.85, *p =* .09]. In addition, analyses of memory for change at the time of test (right panels of **Figure S17**) showed no significant effect of mean PMC reinstatement Z-scores, nor mean PMC reinstatement Z-scores by Age Group interactions (top panels), either between- or within-participants [*largest* χ*^2^*(1) = 2.94, *p =* .09]. However, for the MTL (bottom panels), there were significant Age Group × Reinstatement Z-score interactions both between-participants [χ*^2^*(1) = 4.55, *p* = .03] and within-participants [χ*^2^*(1) = 10.55, *p* = .001]. Young adults with higher mean MTL reinstatement Z-scores remembered significantly more changes [χ*^2^*(1) = 4.83, *p* = .03], but older adults did not [χ*^2^*(1) = 0.04, *p* = .84]. Surprisingly, at the within-participant level, trials with higher reinstatement Z-scores predicted significantly worse memory for change for young adults [χ*^2^*(1) = 5.78, *p* = .02], whereas trials with higher reinstatement Z-scores predicted significantly better memory for change for older adults [χ*^2^*(1) = 4.45, *p* = .03].

Next, for the individual parcel Z-scores, we used logistic mixed effect models with between- and within-participant reinstatement Z-scores but added the individual Z-scores of each parcel as fixed effects rather than a single mean value representing the average of all the parcels (as with the cued recall task, see sections 2.2.2, and 2.2.3). We next used the anova function of the lme4 package (20) to determine whether adding these single parcel Z-scores relative to a baseline model comprising only the random effects improved model fits. We then examined whether these model fits were further improved by adding the interaction terms with Age Group as a fixed effect. Regarding *Day 2 recognition accuracy* for changed activities, model fit was not improved by adding the between-participant reinstatement Z-scores for the PMC or MTL parcels or the interaction terms with Age Group [*largest* χ*^2^*(12) = 10.60, *p =* .56]. However, at the within-participant level, adding the fixed effects of the PMC parcels improved model fit [χ*^2^*(12) = 23.66, *p* = .02], but the interaction terms with Age Group did not [χ*^2^*(12) = 8.76, *p* = .72]. Examination of the parameter estimates for individual parcels showed that higher reinstatement Z-scores in parcel 291 were associated with higher Day 2 recognition accuracy [χ*^2^*(1) = 16.58, *p* < .001]. Adding the MTL parcels at the within-participants level did not improve model fit, nor did the interaction with Age Group [*largest* χ*^2^*(6) = 6.52, *p* = .37].

Regarding memory for change during the recognition task, at the between-participant level, adding the PMC parcels improved model fit [χ*^2^*(12) = 21.52, *p* = .04], as did the interaction with Age Group [χ*^2^*(12) = 21.58, *p* = .04]. Adding the PMC parcels improved model fit in both young [χ*^2^*(12) = 22.91, *p* = .03] and older adults [χ*^2^*(12) = 23.72, *p* = .02], but different individual parcels predicted performance in each age group. In the young adults, higher reinstatement Z-scores in parcels 115 [χ*^2^*(1) = 9.87, *p* = .002] and 291 [χ*^2^*(1) = 6.63, *p* = .01] were associated with better memory for change. In the older adults, higher reinstatement Z-scores in parcel 292 was associated with better memory for change [χ*^2^*(1) = 10.92, *p* < .001], whereas higher reinstatement in parcels 141[χ*^2^*(1) = 8.80, *p* = .003] and 276 [χ*^2^*(1) = 5.29, *p* = .02] predicted worse memory for change. There was no significant effect for the MTL at the between participant level [*largest* χ*^2^*(6) = 10.28, *p* = .11] and no effects for either the PMC or MTL parcels nor the interaction with Age Group were significant when examining the association between reinstatement Z-scores and memory for change at the within-participant level [*largest* χ*^2^*(6) = 11.71, *p =* .07].

Surprisingly, these results do not fully replicate those of the cued recall task at the between-participant level, because mean reinstatement Z-scores were not related to Day 2 recognition accuracy in either age group (although MTL reinstatement Z-scores predicted correctly reporting having seen both versions of the changed activities in the young adults). It is likely that these findings result from the different nature of the two memory tasks. Recall accuracy of the changed Day 2 features relied primarily on recollection processes, whereas recognition of changed features relied relatively less on recollection and more on familiarity. Recent reviews of neuroimaging findings indicate that correct recognition performance does not necessarily rely on activity within posterior default network regions, and that this is particularly the case when participants rely on familiarity rather than recollection processes to guide their memory decisions (40). This lower reliance on PMC and MTL during the recognition task might explain why neural activity pattern reinstatements in these regions did not predict better memory for changed features.

#### d. Hippocampal response for the difference between changed and repeated activities, reinstatement Z-scores, and behavioral performance

To further assess the proposal that hippocampal responses to novelty are a critical component of memory updating when experiencing changes in events (35), we next examined whether the difference in the hippocampal response for changed compared to repeated activities predicted Day 2 recognition of changed activities. To do so, we used linear mixed effect models to examine whether adding these parameter estimates and their interaction with Age Group improved model fits above and beyond reinstatement Z-scores. Results showed that this was the case for Day 2 recognition accuracy for changed items [χ*^2^*(2) = 6.36, *p* = .04, for the MTL; χ*^2^*(2) = 6.43, *p* = .04, for the PMC]. Larger hippocampal response was related to a higher Day 2 recognition accuracy [χ*^2^*(1) = 5.49, *p* = .02, for the MTL; χ*^2^*(1) = 5.49, *p* = .02, for the PMC] and this effect did not interact with Age Group [χ*^2^*(1) = 1.00, *p* = .32, for the MTL; χ*^2^*(1) = 1.07, *p* = .30, for the PMC]. Adding the parameter estimates did not improve model fit for the tendency to remember change in the MTL [χ*^2^*(2) = 3.95, *p* = .14] nor PMC [χ*^2^*(2) = 3.78, *p* = .15].

Together, these results complement those observed in the cued recall task (see section 2.3). Although the intensity of difference in hippocampal responses for changed compared to repeated event viewing does not seem to facilitate subsequent episodic recollection, the present results show that it is nonetheless associated with better Day 2 recognition accuracy for the changed event features. A possible explanation for these discrepant results is that the intensity of hippocampal response has a specific effect on correct recognition based on familiarity processes that are less necessary in the cued recall task. Future studies should investigate whether participants based their answers on recollection or familiarity during similar recognition tasks to assess this possibility.

## 5. Figures

**Figure S1.**
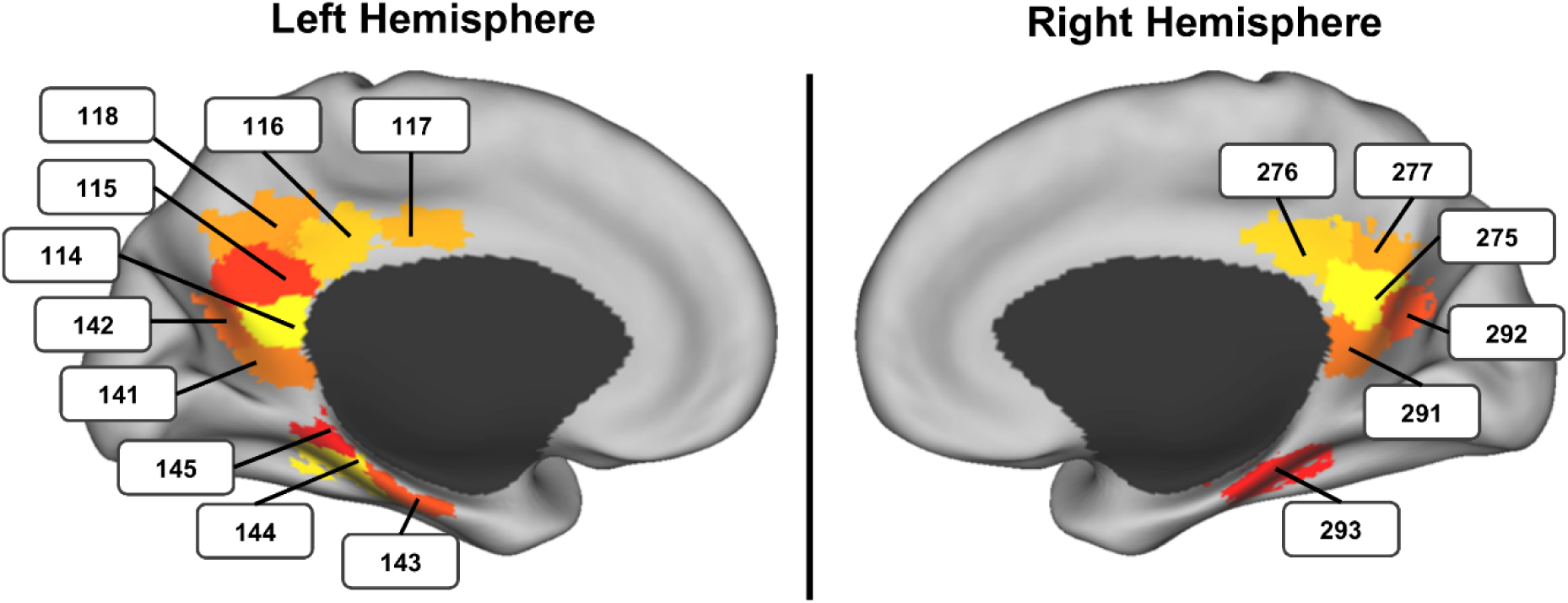
PMC and MTL parcels of interest.

**Figure S2.**
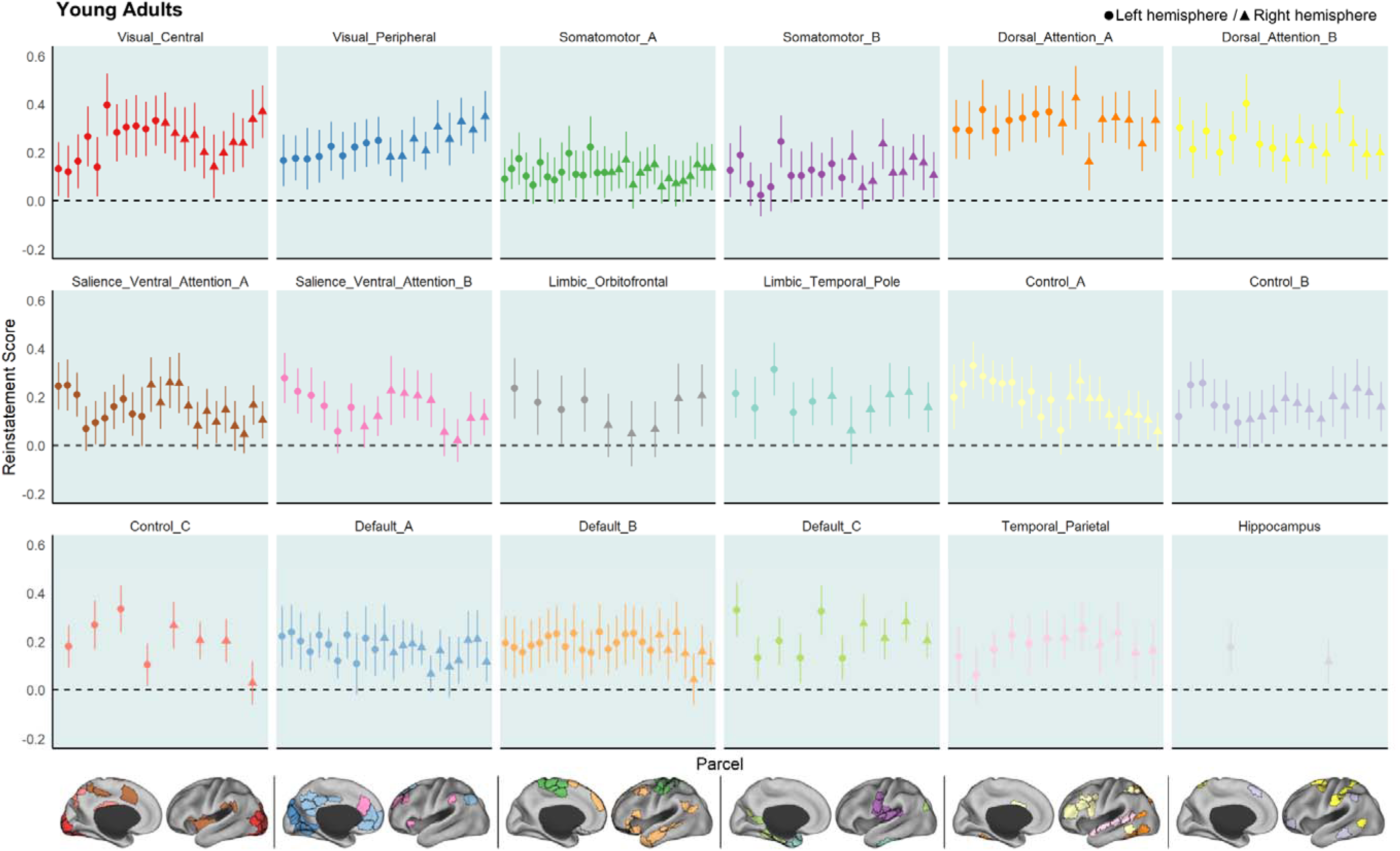
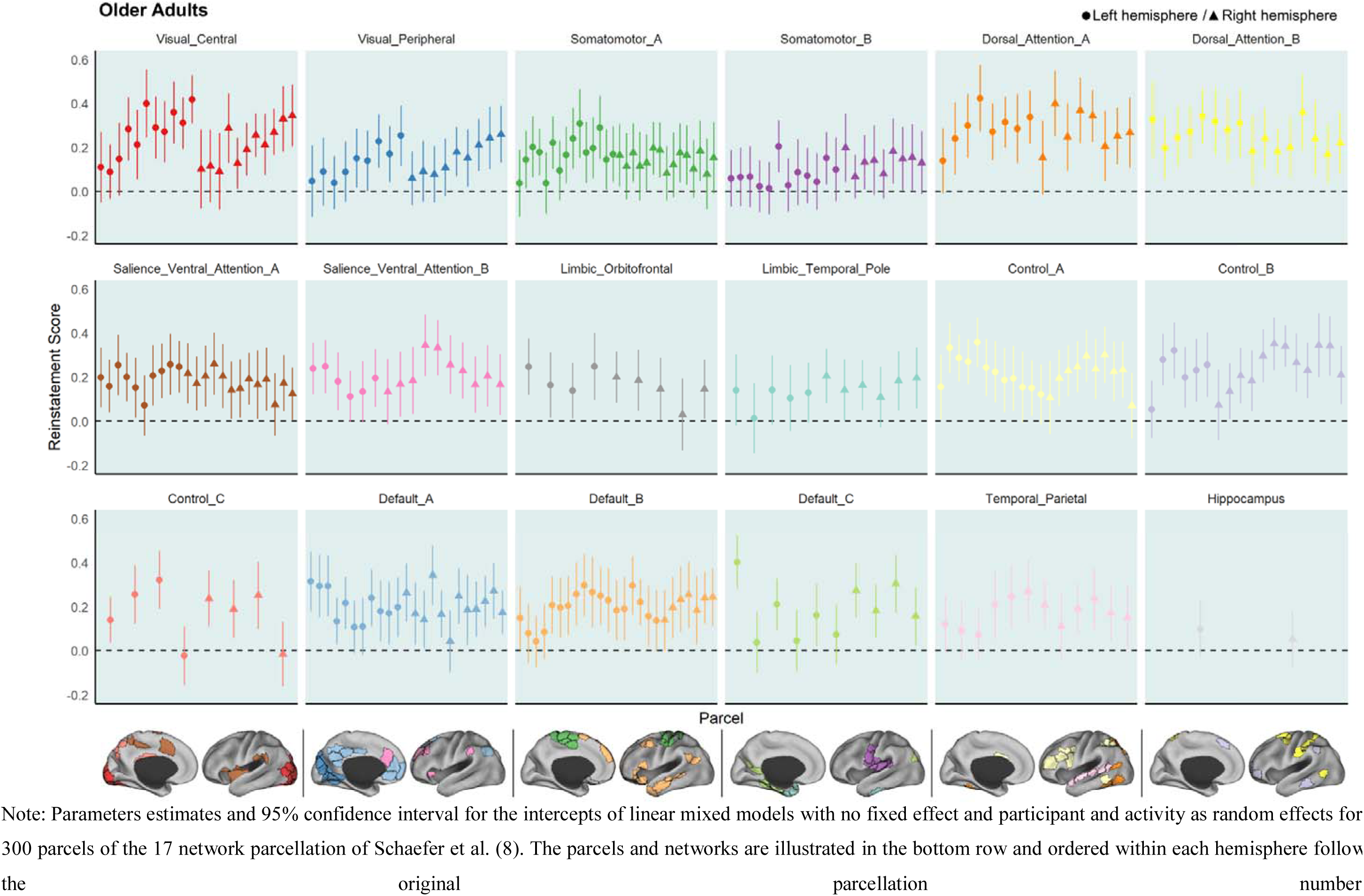
Reinstatement Z-scores across the full set of parcels.

**Figure S3.**
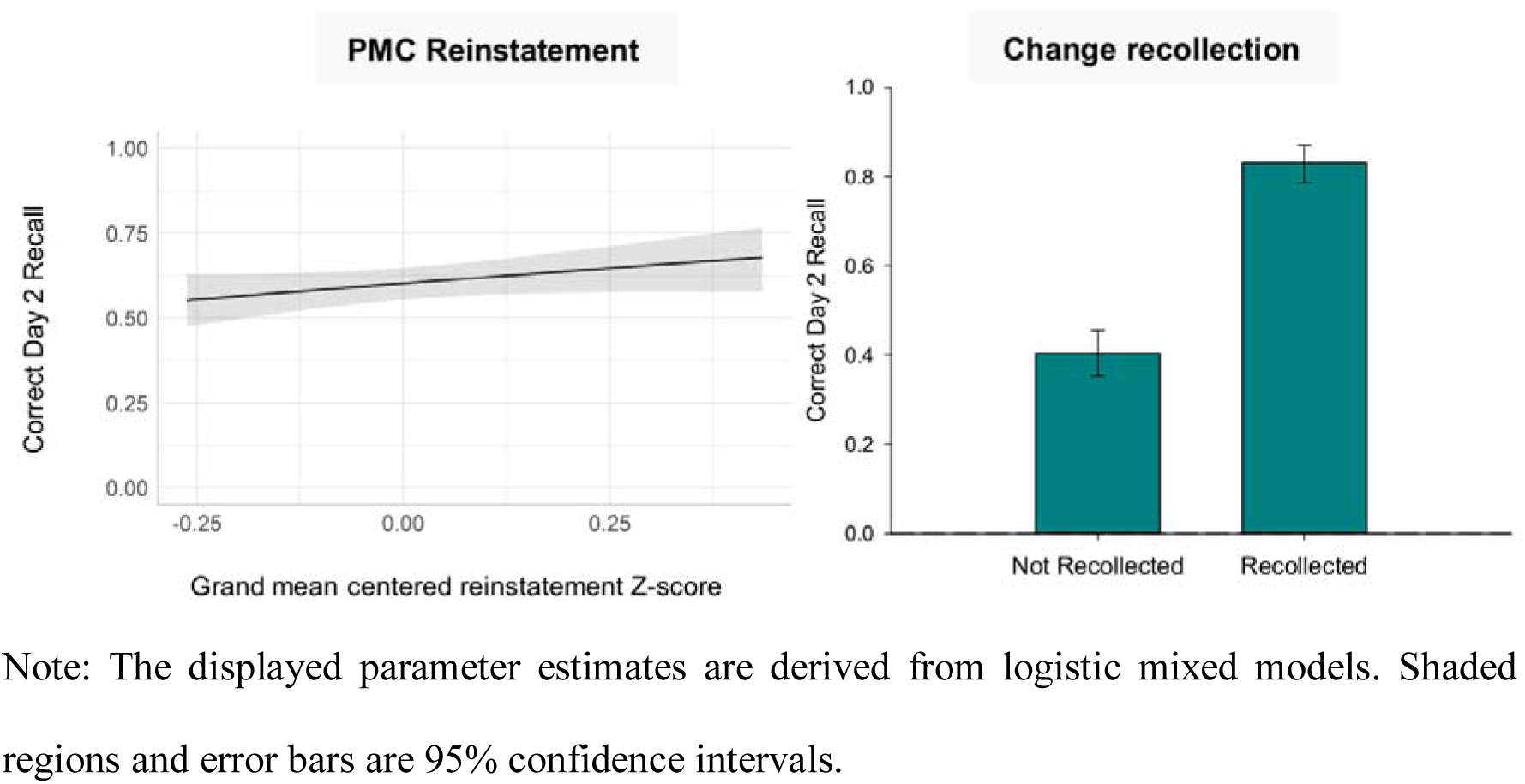
Day 2 recall accuracy by mean PMC/MTL reinstatement Z-scores (between-participant level) and change recollection accuracy for young adults.

**Figure S4.**
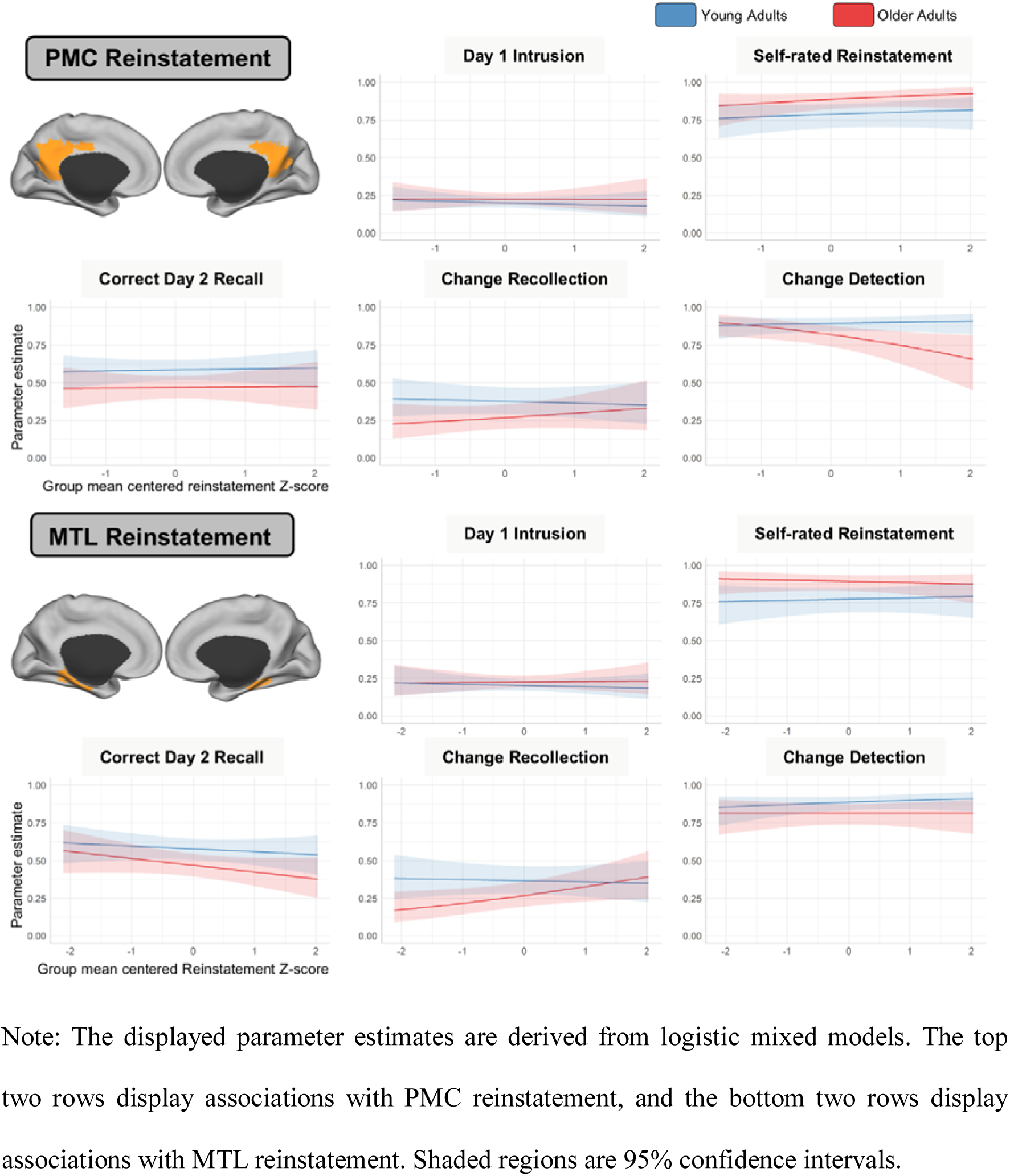
Within-participant associations between neural reinstatement Z-scores and behavioral performance in the cued recall task measures.

**Figure S5.**
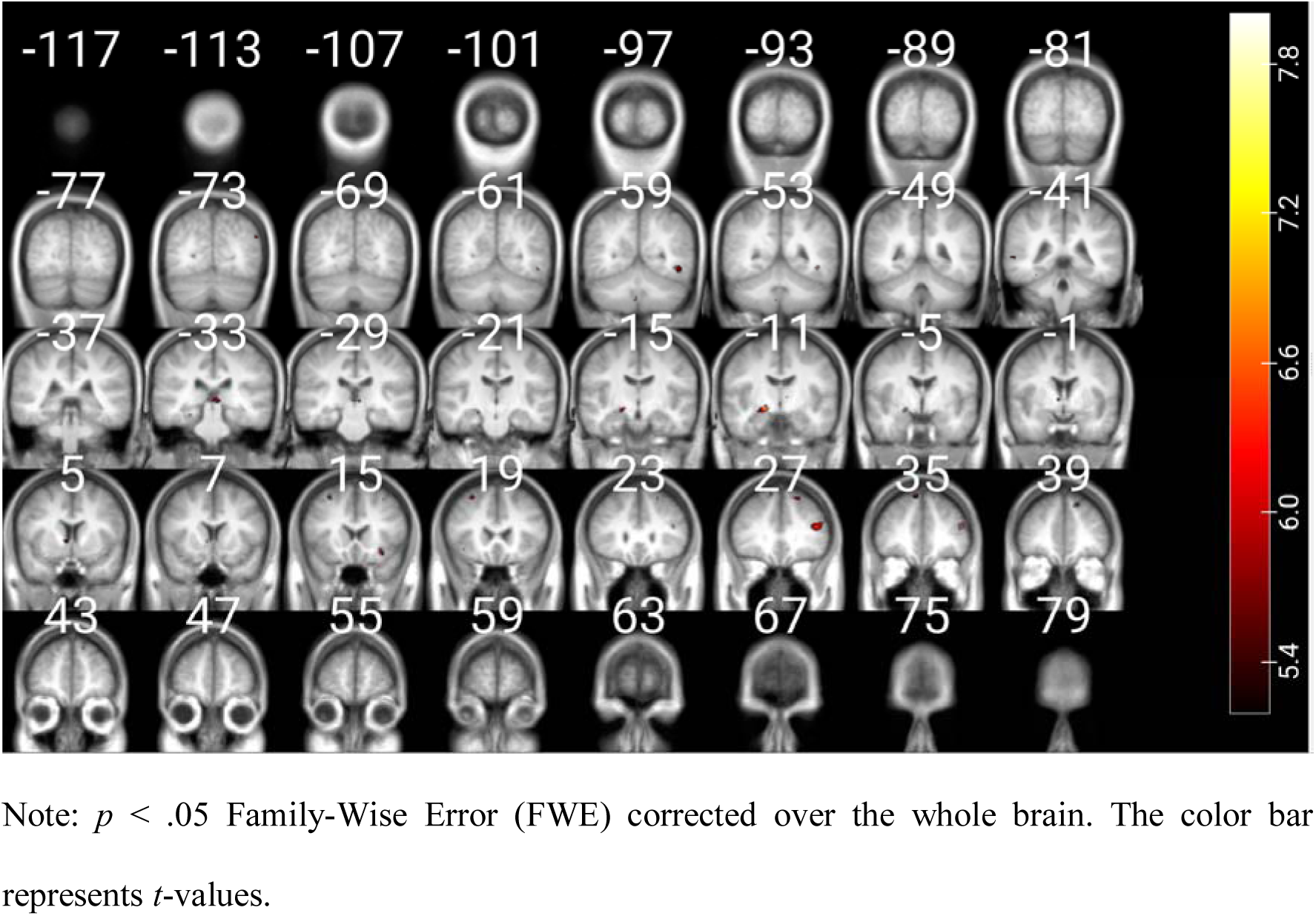
Brain regions more activated while watching changed vs repeated post-divergence segments during the Day 2 viewing across all participants (*N* = 62).

**Figure S6.**
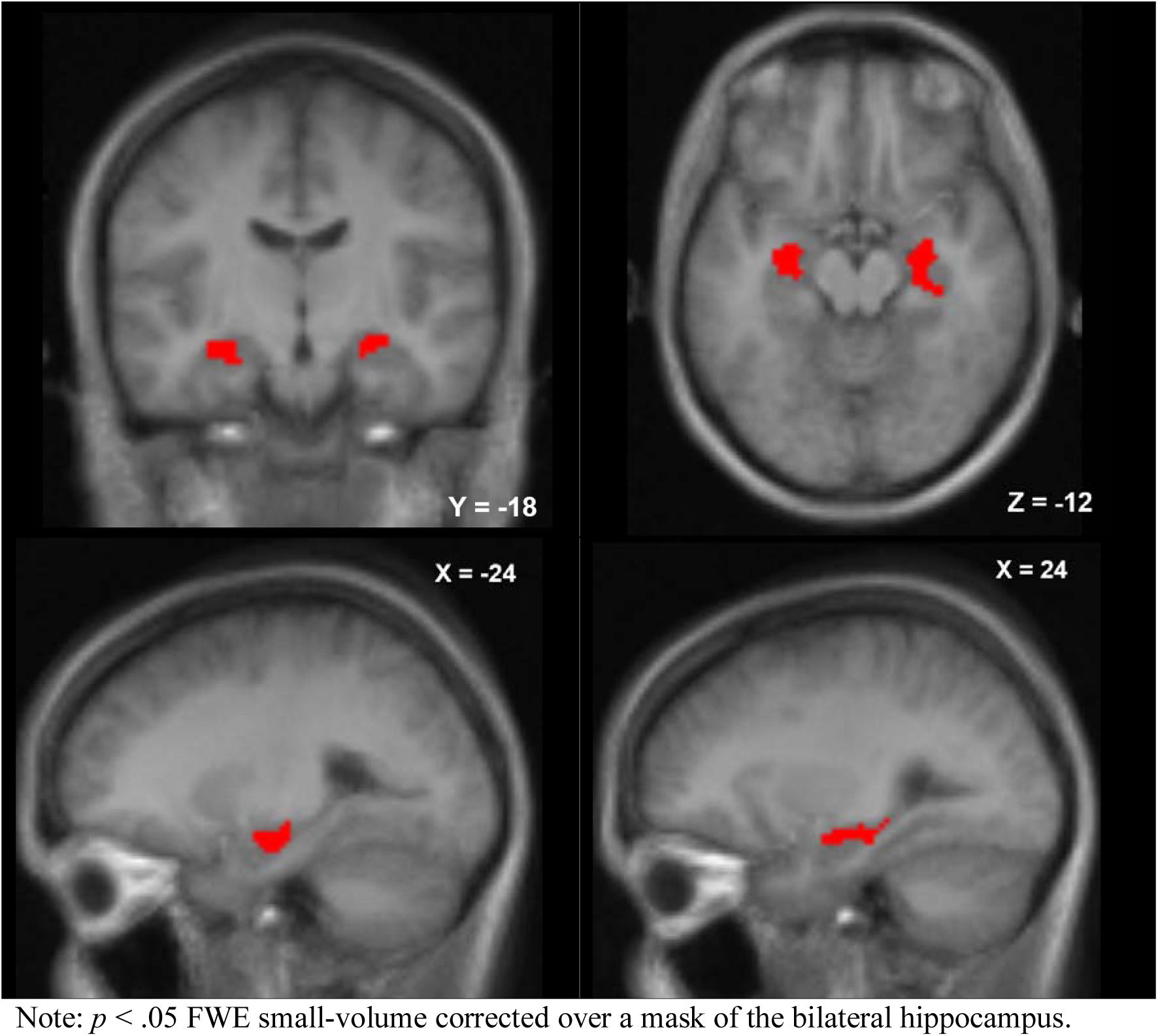
Bilateral anterior hippocampal cluster that is more activated while watching changed than repeated post-divergence segments during Day 2 viewing across all participants (*N* = 62).

**Figure S7.**
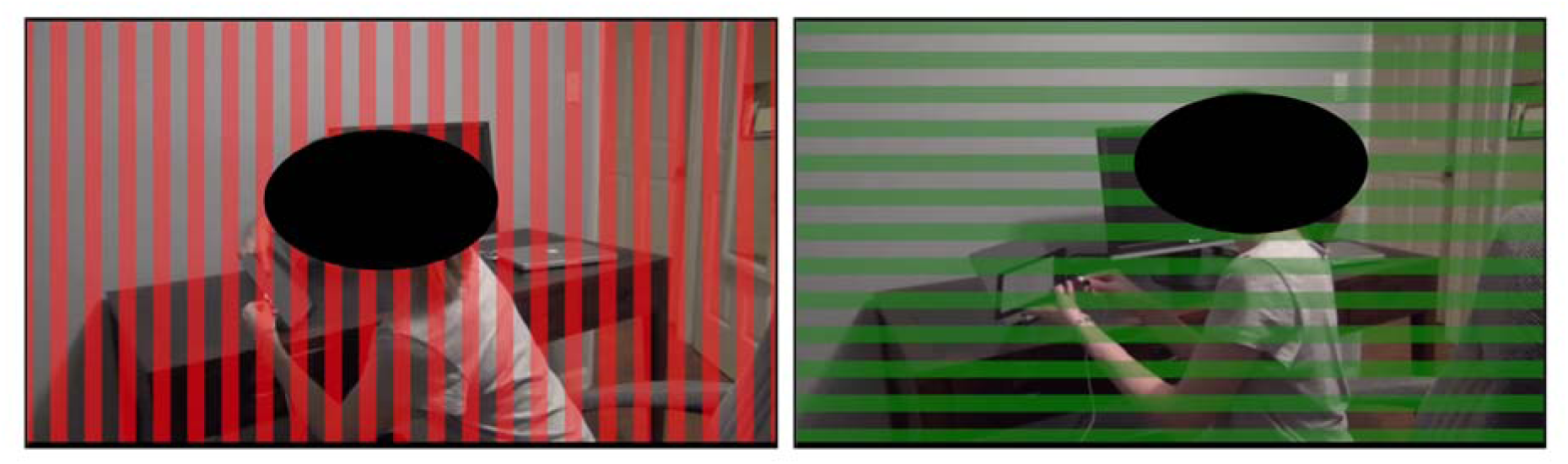
Illustration of the gratings overlayed on the movies during the first 1.5 s of the post-divergence segments of the activities.

**Figure S8.**
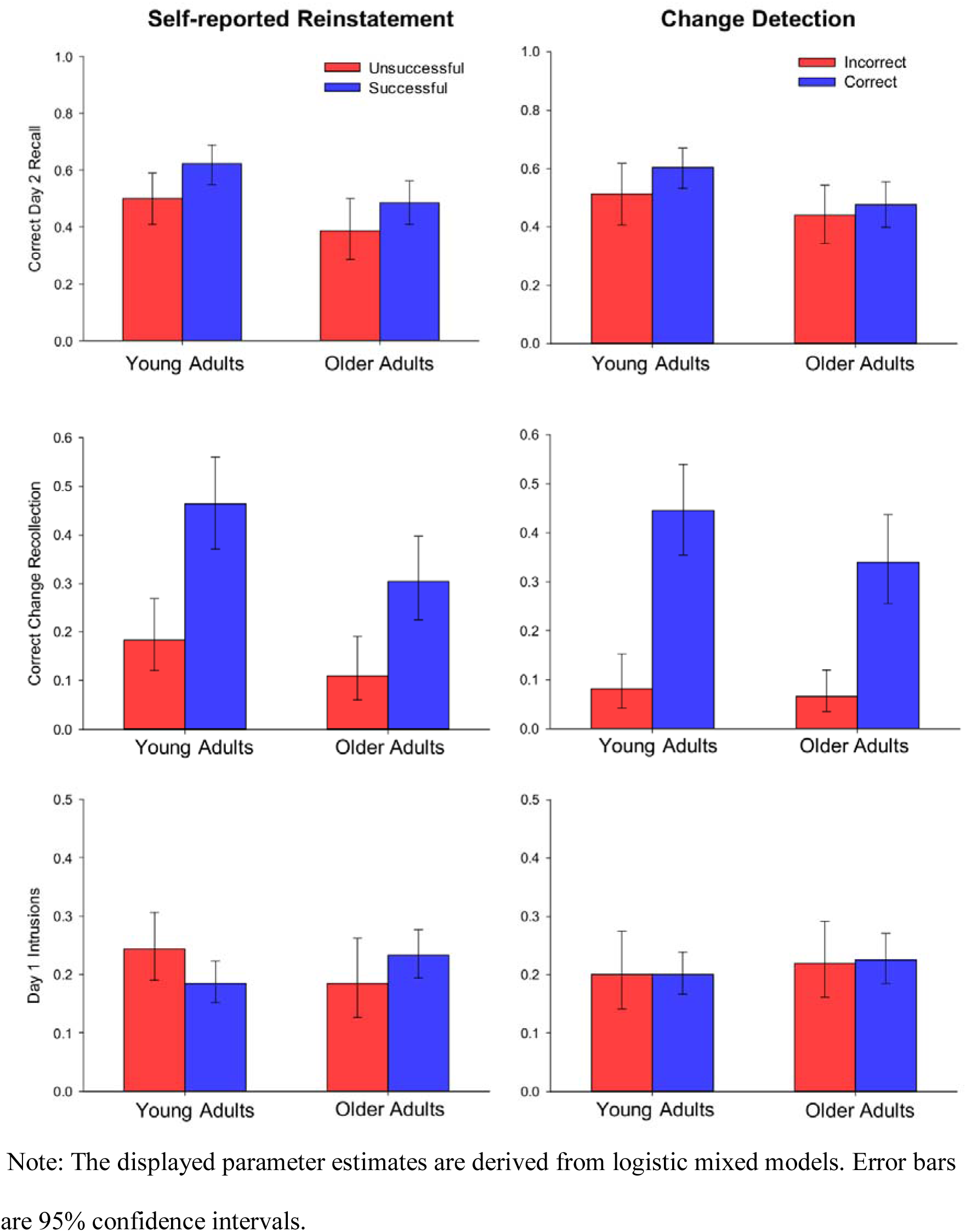
Performance in the cued recall task for changed activities for young and older adults as a function of self-reported reinstatement success and change detection accuracy.

**Figure S9.**
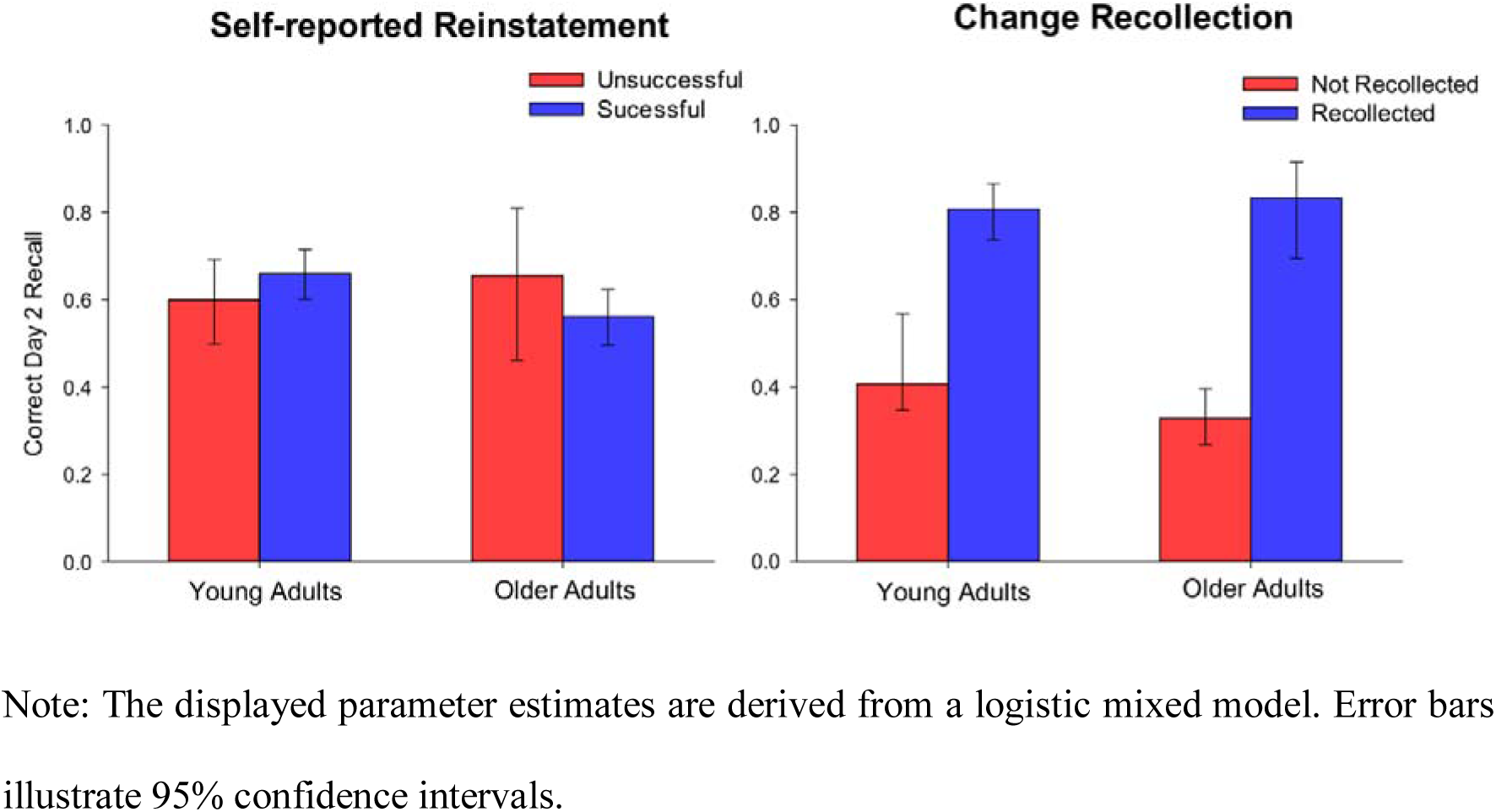
Day 2 recall accuracy conditionalized on self-reported reinstatement success and change recollection accuracy for young and older adults.

**Figure S10.**
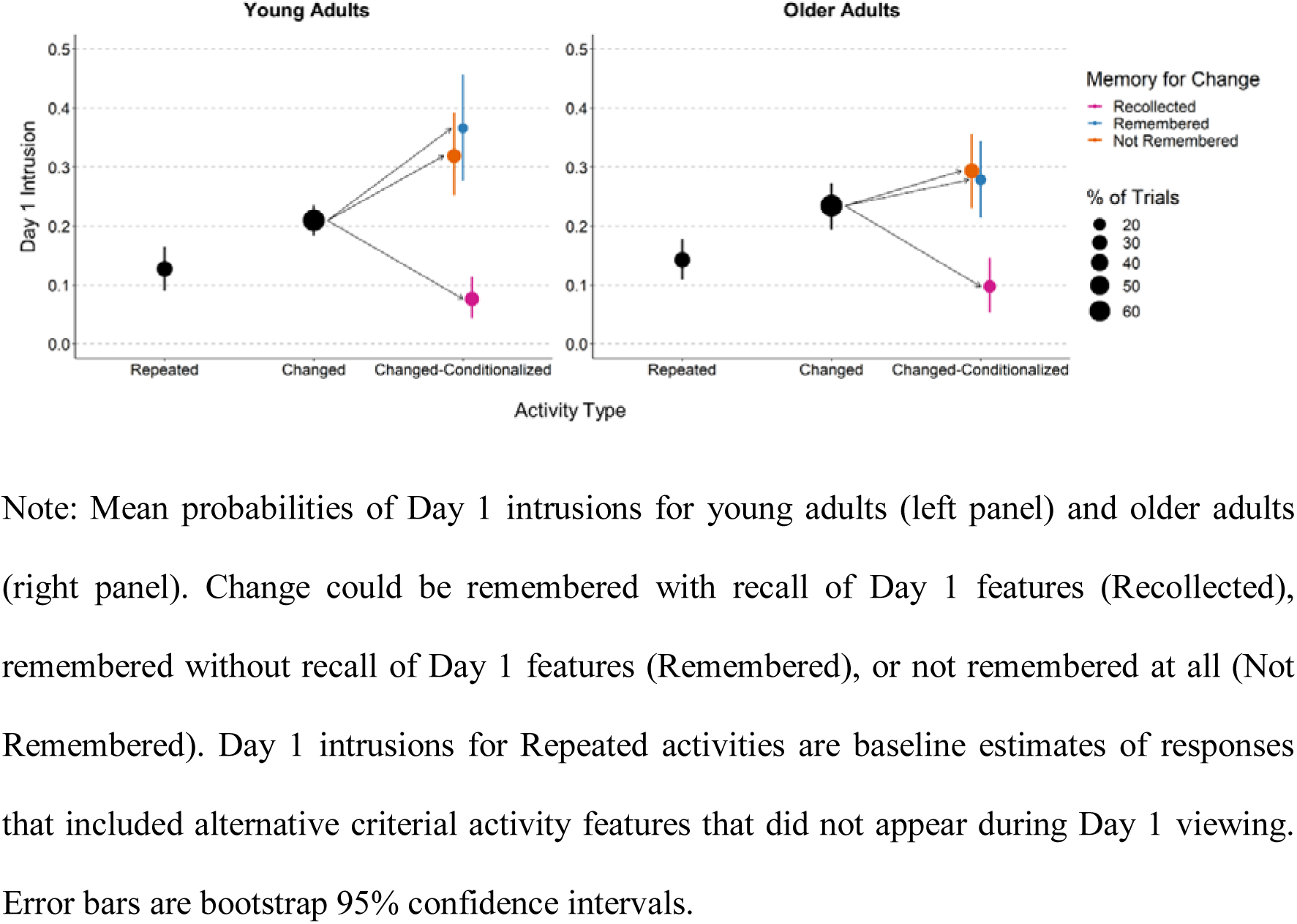
Day 1 Intrusions during Day 2 recall for young and older adults.

**Figure S11.**
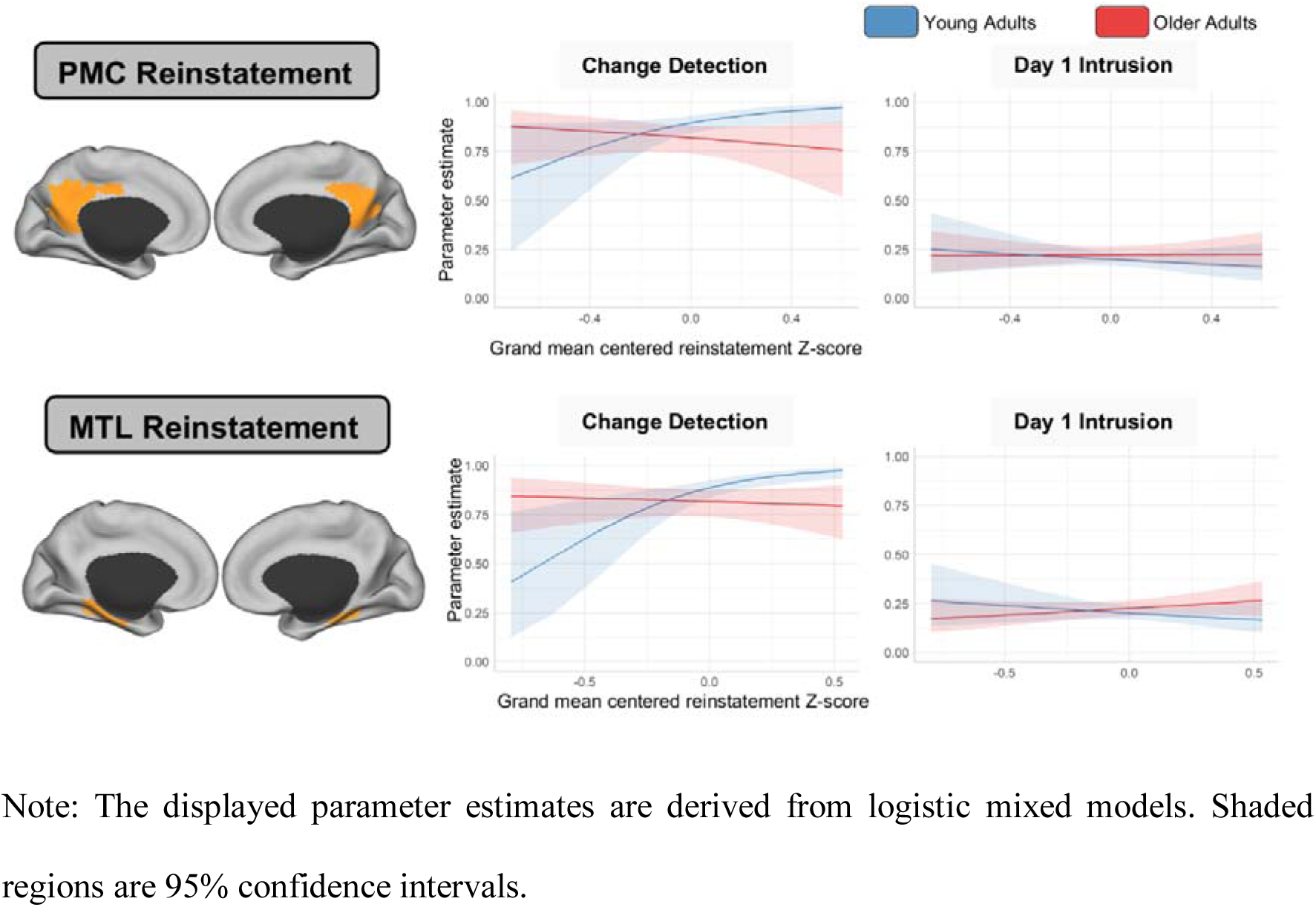
Between-participant associations between behavioral measures (Day 1 Intrusions and Change Detection) and mean PMC/MTL reinstatement Z-scores.

**Figure S12.**
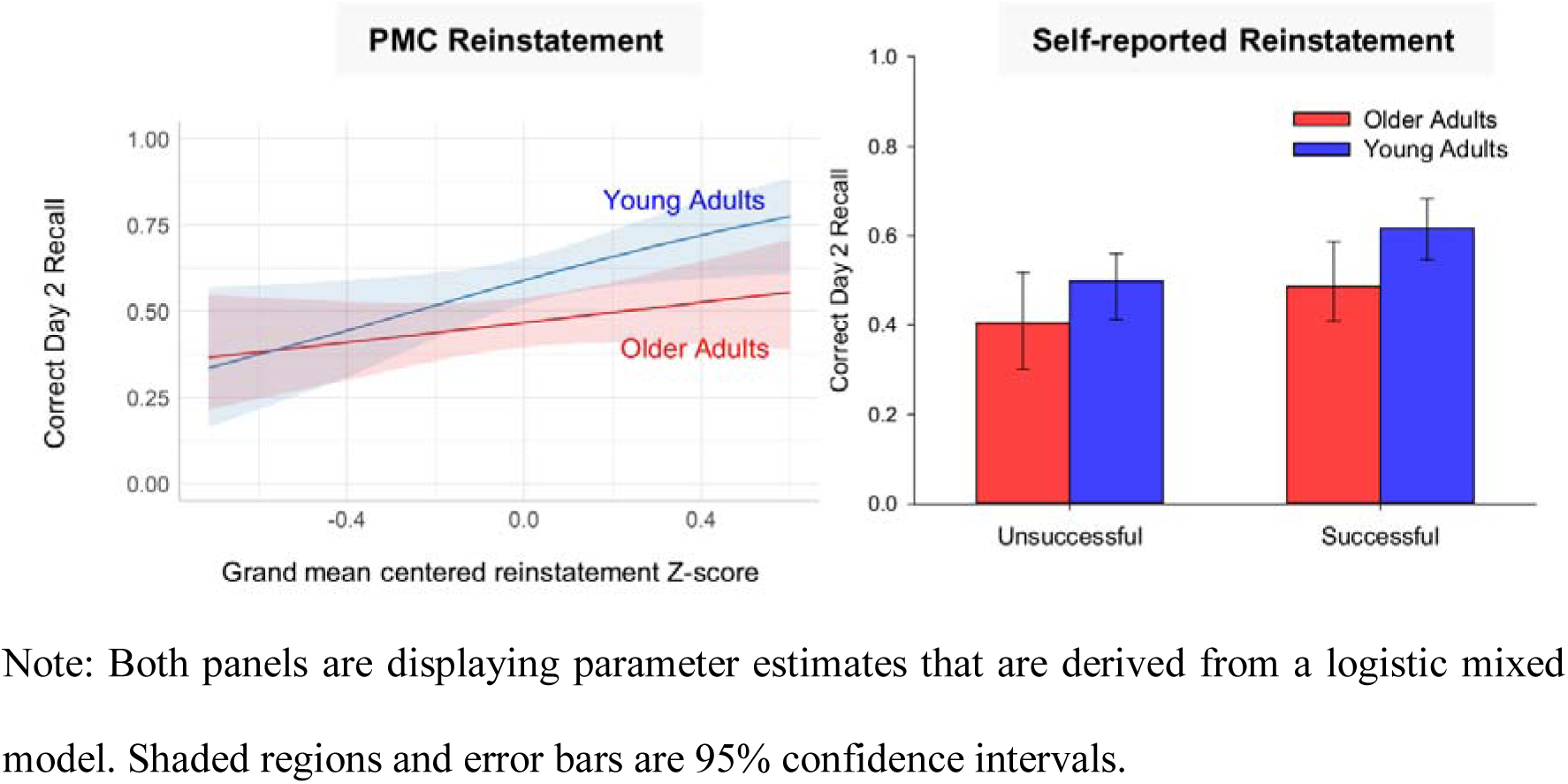
Day 2 recall accuracy by mean PMC reinstatement Z-scores (between-participant level) and self-reported reinstatement success above and beyond the effect of the other variable.

**Figure S13.**
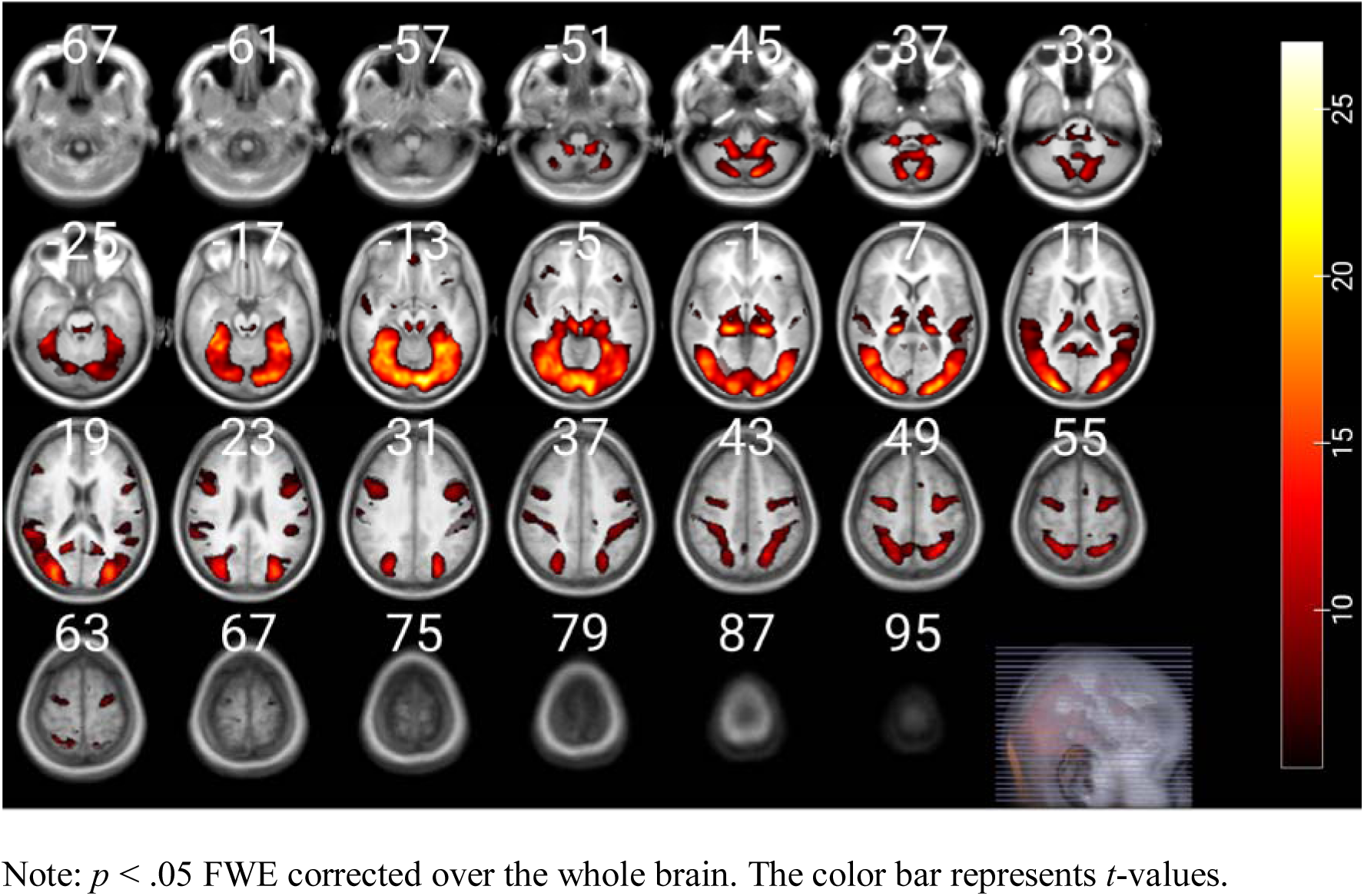
Brain regions that are more activated than baseline while watching activities during Day 1 viewing across all participants (*N* = 62).

**Figure S14.**
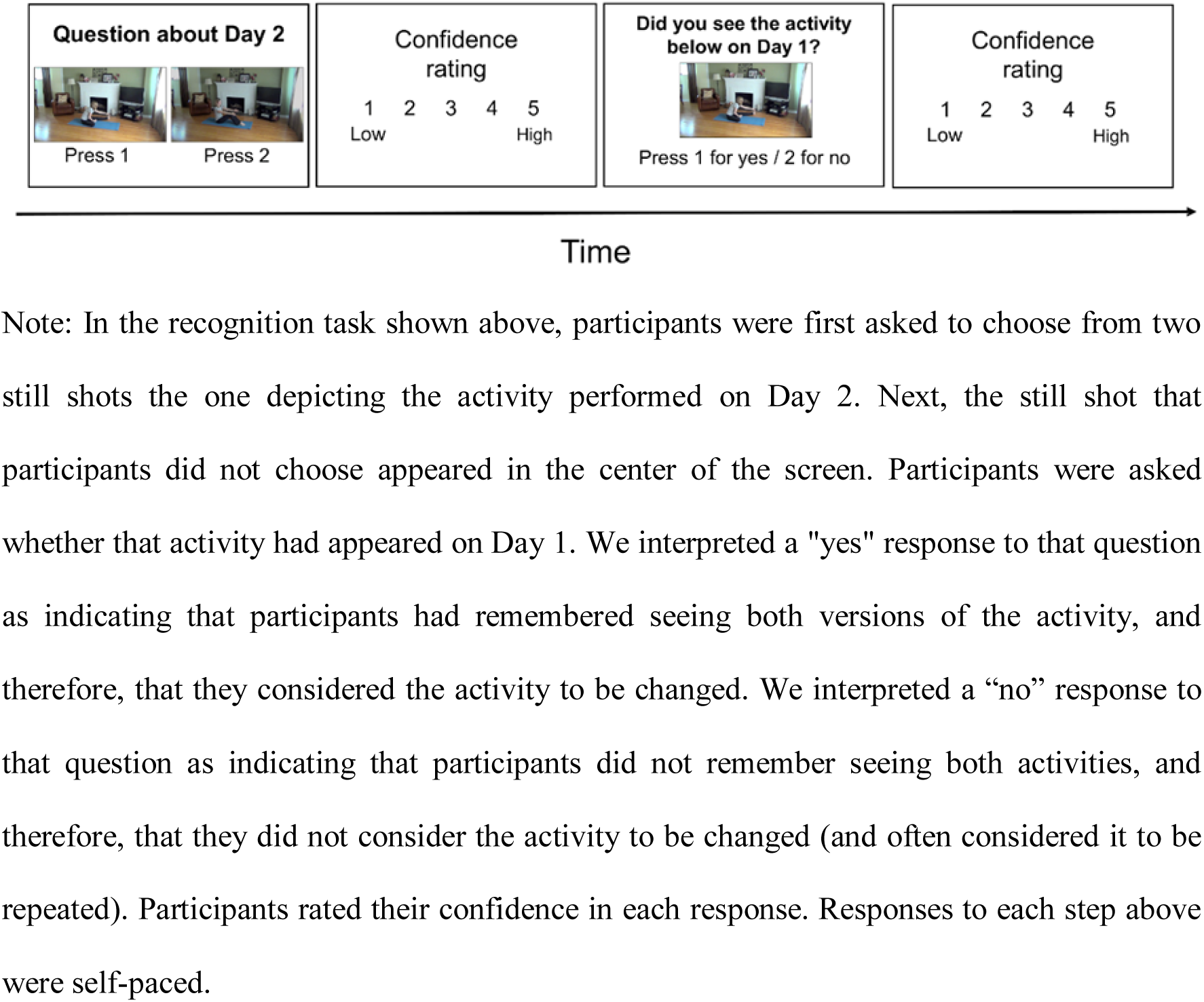
Schematic of the recognition task procedure.

**Figure S15.**
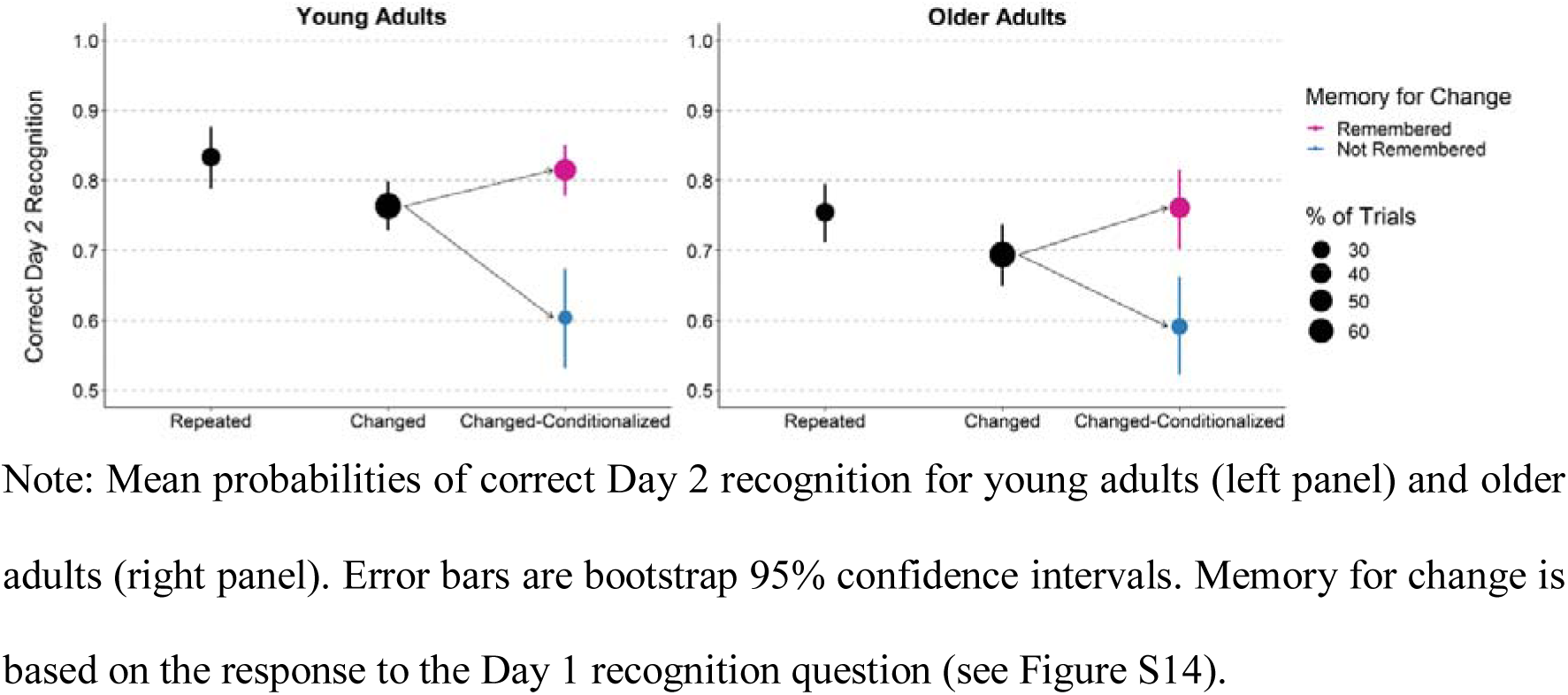
Correct recognition of Day 2 activities for young and older adults.

**Figure S16.**
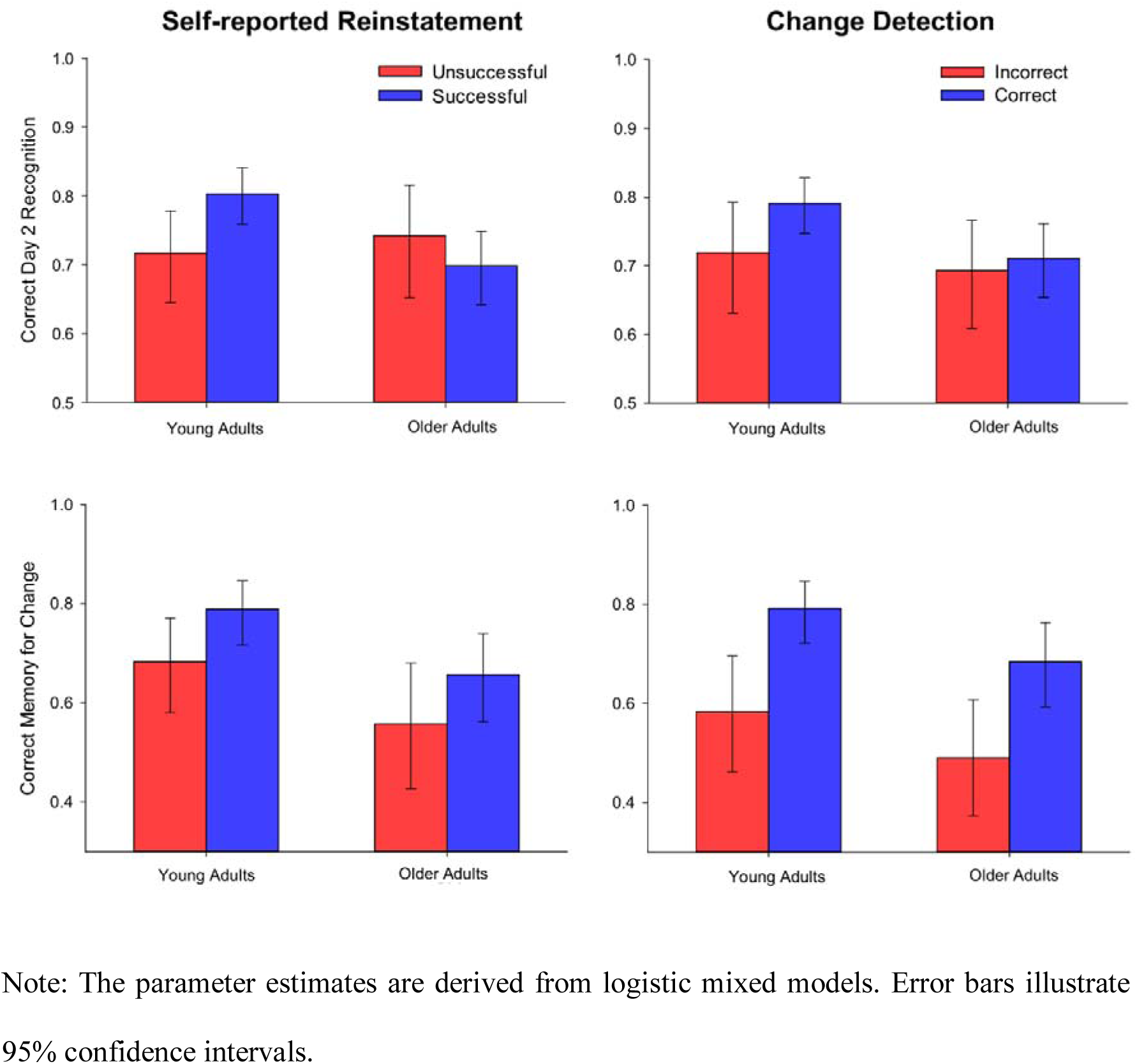
Correct recognition of changed Day 2 activities and correct memory for change in young and older adults conditionalized on self-reported reinstatement success and change detection accuracy.

**Figure S17.**
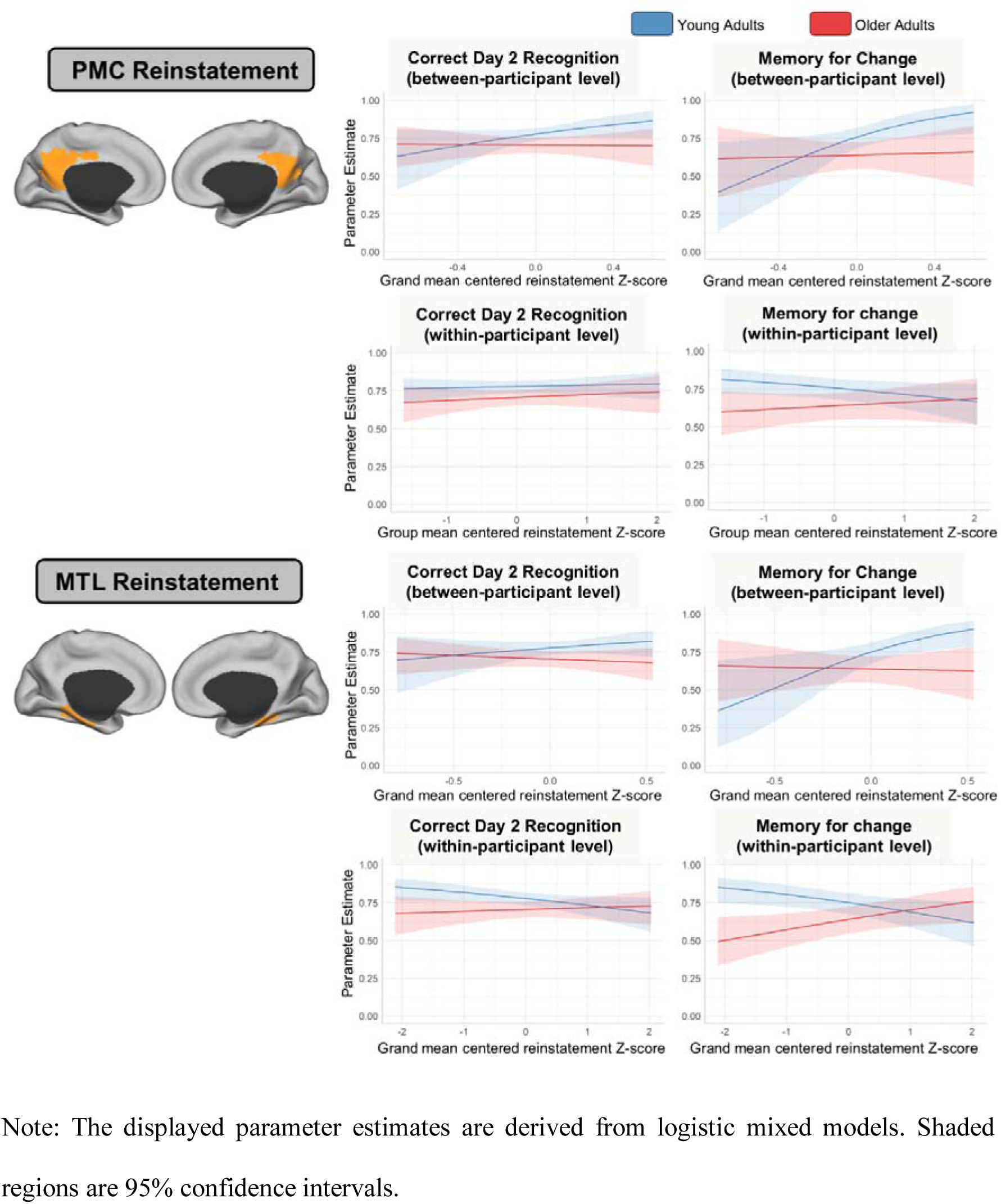
Mean PMC/MTL reinstatement Z-scores predicting recognition memory for changed activities and memory for changes in young and older adults.

## 6. Tables

**Table S1.** Comparisons of reinstatement Z-scores for young and older adults for each PMC and MTL parcel.

| Parcel | $\chi^2(1)$ | *p*-value | Parameter estimates YA<br>[95% CI] | Parameter estimates OA<br>[95% CI] |
| --- | --- | --- | --- | --- |
| **PMC Left** |  |  |  |  |
| p.114 | 0.15 | .70 | 0.16 [0.07; 0.24] | 0.13 [0.04; 0.23] |
| p.115 | 0.02 | .89 | 0.23 [0.125; 0.33] | 0.22 [0.11; 0.33] |
| p.116 | 1.68 | .20 | 0.19 [0.10; 0.27] | 0.11 [0.01; 0.20] |
| p.117 | 0.03 | .87 | 0.12 [0.03; 0.21] | 0.11 [0.01; 0.21] |
| p.118 | 0.03 | .86 | 0.23 [0.12; 0.33] | 0.24 [0.13; 0.35] |
| p.141 | 1.49 | .22 | 0.13 [0.03; 0.235] | 0.04 [-0.08; 0.15] |
| p.142 | 0.01 | .91 | 0.20 [0.10; 0.30] | 0.21 [0.10; 0.32] |
| **PMC Right** |  |  |  |  |
| p.275 | 0.03 | .86 | 0.175 [0.10; 0.25] | 0.165 [0.08; 0.25] |
| p.276 | 0.10 | .75 | 0.07 [-0.03; 0.165] | 0.04 [-0.07; 0.15] |
| p.277 | 1.98 | .16 | 0.16 [0.07; 0.26] | 0.25 [0.14; 0.35] |
| p.291 | 0.24 | .63 | 0.21 [0.12; 0.31] | 0.18 [0.08; 0.28] |
| p.292 | 0.10 | .75 | 0.28 [0.18; 0.38] | 0.305 [0.20; 0.41] |
| **MTL Left** |  |  |  |  |
| p.143 | 1.11 | .29 | 0.13 [0.02; 0.24] | 0.04 [-0.08; 0.17] |
| p.144 | 3.94 | .047 | 0.33 [0.21; 0.44] | 0.16 [0.04; 0.29] |
| p.145 | 0.58 | .44 | 0.13 [0.03; 0.24] | 0.07 [-0.04; 0.19] |
| Hippo.left | 1.02 | .31 | 0.18 [0.07; 0.28] | 0.10 [-0.02; 0.21] |
| **MTL Right** |  |  |  |  |
| p.293 | 0.46 | .50 | 0.20 [0.11; 0.30] | 0.5 [-0.06; 0.16] |
| Hippo.Right | 0.76 | .38 | 0.12 [0.02; 0.22] | 0.01 [-0.01; 0.04] |

**Table S2.** Pearson correlation matrix of reinstatement Z-scores.

|  | 114 | 115 | 116 | 117 | 118 | 141 | 142 | 277 | 275 | 276 | 291 | 292 | 143 | 144 | 145 | Hip.L | 293 |
| --- | --- | --- | --- | --- | --- | --- | --- | --- | --- | --- | --- | --- | --- | --- | --- | --- | --- |
| 115 | .37 |  |  |  |  |  |  |  |  |  |  |  |  |  |  |  |  |
| 116 | .23 | .33 |  |  |  |  |  |  |  |  |  |  |  |  |  |  |  |
| 117 | .14 | .18 | .25 |  |  |  |  |  |  |  |  |  |  |  |  |  |  |
| 118 | .26 | .45 | .36 | .20 |  |  |  |  |  |  |  |  |  |  |  |  |  |
| 141 | .30 | .25 | .20 | .10 | .24 |  |  |  |  |  |  |  |  |  |  |  |  |
| 142 | .27 | .33 | .19 | .10 | .29 | .29 |  |  |  |  |  |  |  |  |  |  |  |
| 277 | .36 | .43 | .25 | .16 | .30 | .22 | .25 |  |  |  |  |  |  |  |  |  |  |
| 275 | .25 | .31 | .43 | .24 | .31 | .19 | .20 | .31 |  |  |  |  |  |  |  |  |  |
| 276 | .24 | .38 | .26 | .17 | .40 | .20 | .23 | .40 | .33 |  |  |  |  |  |  |  |  |
| 291 | .30 | .29 | .24 | .15 | .27 | .34 | .26 | .33 | .27 | .26 |  |  |  |  |  |  |  |
| 292 | .21 | .24 | .15 | .09 | .24 | .25 | .34 | .30 | .22 | .27 | .34 |  |  |  |  |  |  |
| 143 | .13 | .14 | .17 | .15 | .15 | .18 | .12 | .10 | .16 | .12 | .17 | .11 |  |  |  |  |  |
| 144 | .17 | .21 | .23 | .11 | .23 | .26 | .24 | .15 | .20 | .18 | .24 | .21 | .32 |  |  |  |  |
| 145 | .15 | .17 | .18 | .11 | .16 | .24 | .14 | .13 | .18 | .16 | .22 | .16 | .25 | .32 |  |  |  |
| Hip.L | .16 | .15 | .16 | .14 | .16 | .17 | .15 | .15 | .16 | .12 | .18 | .12 | .34 | .39 | .23 |  |  |
| 293 | .18 | .21 | .17 | .13 | .19 | .19 | .19 | .18 | .23 | .19 | .29 | .22 | .28 | .33 | .20 | .23 |  |
| Hip.R | .13 | .15 | .14 | .12 | .16 | .13 | .12 | .17 | .21 | .14 | .21 | .15 | .20 | .21 | .12 | .27 | .45 |

Note: The coefficients were obtained by correlating the reinstatement Z-scores of each trial across all participants.

**Table S3:** Self-reported reinstatement success (reinstatement success) and activity type classification accuracy (classification accuracy) during Day 2 viewing based on activity type and age group.

| Measure | Age Group | Activity Type |  |
| --- | --- | --- | --- |
|  |  | Repeated | Changed |
| **Reinstatement Success** | Young | .80 [.71, .87] | .79 [.70, .86] |
|  | Older | .90 [.83, .94] | .88 [.81, .93] |
| **Classification Accuracy** | Young | .87 [.81, .91] | .88 [.83, .91] |
|  | Older | .79 [.71, .85] | .80 [.73, .85] |

Note: The displayed parameter estimates are derived from logistic mixed models. 95% confidence intervals are displayed in brackets.

**Table S4:**
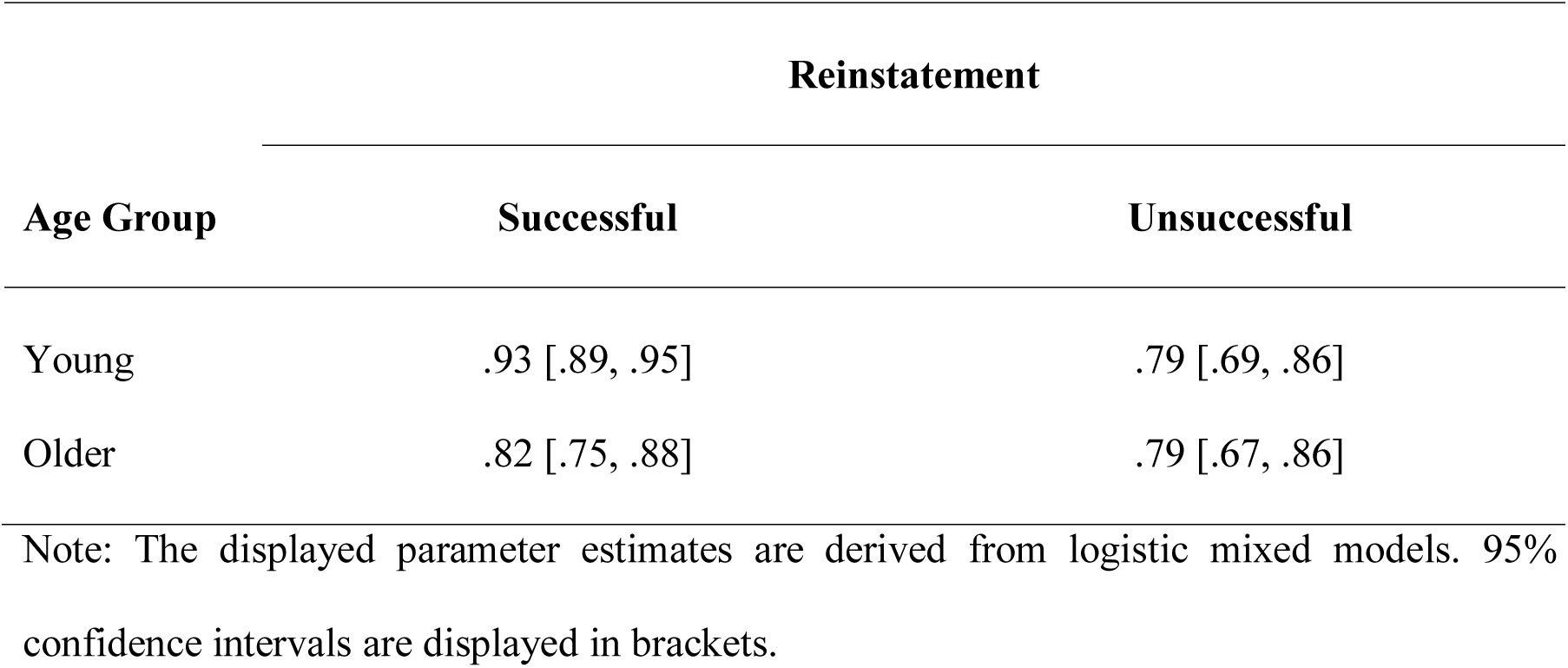
Classification accuracy for changed activities during Day 2 viewing (change detection) conditionalized on self-reported reinstatement success (reinstatement) for both age groups.

| Age Group | Reinstatement |  |
| --- | --- | --- |
|  | Successful | Unsuccessful |
| Young | .93 [.89, .95] | .79 [.69, .86] |
| Older | .82 [.75, .88] | .79 [.67, .86] |

Note: The displayed parameter estimates are derived from logistic mixed models. 95% confidence intervals are displayed in brackets.

## Footnotes

1 The aim of these gratings was to provide an alternative means of analyzing neural pattern reinstatement that allowed for repeated presentations of stimulus features as a complement to identifying unique event features. Initial analyses indicated that reinstatement of event features was more robust than reinstatement of grating features, so the effects of the gratings were not analyzed further.

2 Given the lack of difference in Day 2 recall performance when change was not remembered or remembered without the Day 1 feature (see Figure 2), we collapsed across these cells in this and subsequent analyses. Conditional cells for changed activities were classified as *change recollected* (remembered as changed and the Day 1 feature correctly recalled) and *change not recollected* (not remembered as changed or remembered without the Day 1 feature).

## References

1. D. C. Rubin, S. Umanath, Event memory: A theory of memory for laboratory, autobiographical, and fictional events. Psychol Rev 122, 1–23 (2015).

2. E. Tulving, Episodic memory: from mind to brain. Annu Rev Psychol 53, 1–25 (2002).

3. M. Bar, E. Aminoff, M. F. Mason, M. Fenske, The units of thought. Hippocampus 17, 420–428 (2007).

4. D. L. Schacter, et al., The future of memory: remembering, imagining, and the brain. Neuron 76, 677–94 (2012).

5. C. N. Wahlheim, J. M. Zacks, Memory guides the processing of event changes for older and younger adults. Journal of Experimental Psychology: General 148, 30–50 (2019).

6. L. Nadel, A. Hupbach, R. Gomez, K. Newman-Smith, Memory formation, consolidation and transformation. Neuroscience & Biobehavioral Reviews 36, 1640–1645 (2012).

7. M. L. Schlichting, A. R. Preston, “The Hippocampus and Memory Integration: Building Knowledge to Navigate Future Decisions” in The Hippocampus from Cells to Systems: Structure, Connectivity, and Functional Contributions to Memory and Flexible Cognition, D. E. Hannula, M. C. Duff, Eds. (Springer International Publishing, 2017), pp. 405–437.

8. D. Clewett, L. Davachi, The ebb and flow of experience determines the temporal structure of memory. Current Opinion in Behavioral Sciences 17, 186–193 (2017).

9. F. R. Richter, A. J. H. Chanales, B. A. Kuhl, Predicting the integration of overlapping memories by decoding mnemonic processing states during learning. NeuroImage 124, 323–335 (2016).

10. K. A. Norman, E. L. Newman, G. Detre, A neural network model of retrieval-induced forgetting. Psychological Review 114, 887–953 (2007).

11. G. Kim, J. A. Lewis-Peacock, K. A. Norman, N. B. Turk-Browne, Pruning of memories by context-based prediction error. Proceedings of the National Academy of Sciences 111, 8997–9002 (2014).

12. S. DuBrow, L. Davachi, Temporal memory is shaped by encoding stability and intervening item reactivation. Journal of Neuroscience 34, 13998–14005 (2014).

13. K. L. Bottenhorn, et al., Cooperating yet distinct brain networks engaged during naturalistic paradigms: A meta-analysis of functional MRI results. Network Neuroscience 3, 27–48 (2018).

14. D. A. Balota, P. O. Dolan, J. M. Duchek, “Memory changes in healthy older adults” in (Oxford University Press, 2000), pp. 395–410.

15. S. R. Old, M. Naveh-Benjamin, Differential effects of age on item and associative measures of memory: A meta-analysis. Psychology and Aging 23, 104–118 (2008).

16. J. F. Danker, J. R. Anderson, The Ghosts of Brain States Past: Remembering Reactivates the Brain Regions Engaged During Encoding. Psychological Bulletin 136, 87–102 (2010).

17. M. E. Wheeler, S. E. Petersen, R. L. Buckner, Memory’s echo: vivid remembering reactivates sensory-specific cortex.[Erratum appears in Proc Natl Acad Sci U S A. 2004 Apr 6;101(14):5181]. Proc Natl Acad Sci U S A 97, 11125–9 (2000).

18. J. Chen, et al., Shared memories reveal shared structure in neural activity across individuals. Nature Neuroscience 20, 115–125 (2017).

19. C. M. Bird, J. L. Keidel, L. P. Ing, A. J. Horner, N. Burgess, Consolidation of Complex Events via Reinstatement in Posterior Cingulate Cortex. J Neurosci 35, 14426–34 (2015).

20. M. E. Raichle, et al., A default mode of brain function. Proc Natl Acad Sci U S A 98, 676–82 (2001).

21. C. Ranganath, M. Ritchey, Two cortical systems for memory-guided behaviour. Nature Reviews Neuroscience 13, 713–726 (2012).

22. A. W. Gilmore, S. M. Nelson, K. B. McDermott, The Contextual Association Network Activates More for Remembered than for Imagined Events. Cereb Cortex 26, 611–7 (2016).

23. D. Stawarczyk, O. Jeunehomme, A. D’Argembeau, Differential contributions of default and dorsal Attention networks to remembering thoughts and external stimuli from real-life events. Cereb Cortex 28, 4023–4035 (2018).

24. N. Raz, K. M. Rodrigue, Differential aging of the brain: Patterns, cognitive correlates and modifiers. Neuroscience & Biobehavioral Reviews 30, 730–748 (2006).

25. R. Leech, D. J. Sharp, The role of the posterior cingulate cortex in cognition and disease. Brain 137, 12–32 (2014).

26. F. Lin, et al., The cingulate cortex of older adults with excellent memory capacity. Cortex 86, 83–92 (2017).

27. N. Kriegeskorte, M. Mur, P. A. Bandettini, Representational similarity analysis - connecting the branches of systems neuroscience. Front. Syst. Neurosci. 2 (2008).

28. A. Schaefer, et al., Local-Global Parcellation of the Human Cerebral Cortex from Intrinsic Functional Connectivity MRI. Cereb Cortex 28, 3095–3114 (2018).

29. S. E. Favila, A. J. H. Chanales, B. A. Kuhl, Experience-dependent hippocampal pattern differentiation prevents interference during subsequent learning. Nature Communications 7 (2016).

30. M. D. Fox, et al., The human brain is intrinsically organized into dynamic, anticorrelated functional networks. PNAS Proceedings of the National Academy of Sciences of the United States of America 102, 9673–9678 (2005).

31. D. Stawarczyk, M. A. Bezdek, J. M. Zacks, Event Representations and Predictive Processing: The Role of the Midline Default Network Core. Topics in Cognitive Science (In press) https:/doi.org/10.1111/tops.12450 (September 6, 2019).

32. C. Baldassano, et al., Discovering Event Structure in Continuous Narrative Perception and Memory. Neuron 95, 709–721.e5 (2017).

33. R. K. Olsen, S. N. Moses, L. Riggs, J. D. Ryan, The hippocampus supports multiple cognitive processes through relational binding and comparison. Front. Hum. Neurosci. 6 (2012).

34. A. Mayes, D. Montaldi, E. Migo, Associative memory and the medial temporal lobes. Trends in Cognitive Sciences 11, 126–135 (2007).

35. A. Ben-Yakov, R. N. Henson, The Hippocampal Film Editor: Sensitivity and Specificity to Event Boundaries in Continuous Experience. J. Neurosci. 38, 10057–10068 (2018).

36. B. Levine, E. Svoboda, J. F. Hay, G. Winocur, M. Moscovitch, Aging and autobiographical memory: dissociating episodic from semantic retrieval. Psychol Aging 17, 677–689 (2002).

37. C. A. Kurby, J. M. Zacks, Age differences in the perception of hierarchical structure in events. Mem Cognit 39, 75–91 (2011).

38. A. Gazzaley, M. D’Esposito, BOLD functional MRI and cognitivec aging (Oxford University Press., 2005).

39. J. L. Earles, A. Kersten, B. Berlin Mas, D. M. Miccio, Aging and Memory for Self-Performed Tasks: Effects of Task Difficulty and Time Pressure. J Gerontol B Psychol Sci Soc Sci 59, P285–P293 (2004).

40. J. J. Allaire, et al., rmarkdown: Dynamic Documents for R. R package version 1.13 (2019).

41. M. F. Folstein, S. E. Folstein, P. R. McHugh, “Mini-mental state”. A practical method for grading the cognitive state of patients for the clinician. J Psychiatr Res 12, 189–198 (1975).

42. W. C. Shipley, C. C. Burlingame, A convenient self-administering scale for measuring intellectual impairment in psychotics. AJP 97, 1313–1325 (1941).

## References

1. W. C. Shipley, C. C. Burlingame, A convenient self-administering scale for measuring intellectual impairment in psychotics. AJP 97, 1313–1325 (1941).

2. W. Schneider, A. Eschman, A. Zuccolotto, E-Prime user’s guide (Psychology Software Tools Inc., 2012).

3. M. F. Folstein, S. E. Folstein, P. R. McHugh, “Mini-mental state”. A practical method for grading the cognitive state of patients for the clinician. J Psychiatr Res 12, 189–198 (1975).

4. R. L. Buckner, et al., A unified approach for morphometric and functional data analysis in young, old, and demented adults using automated atlas-based head size normalization: reliability and validation against manual measurement of total intracranial volume. NeuroImage 23, 724–738 (2004).

5. J. L. R. Andersson, M. Jenkinson, S. Smith, Non-linear registration, aka spatial normalisation. FMRIB technical report TR07JA2 (2010).

6. M. Jenkinson, C. F. Beckmann, T. E. J. Behrens, M. W. Woolrich, S. M. Smith, FSL. NeuroImage 62, 782–790 (2012).

7. J. L. R. Andersson, S. Skare, J. Ashburner, How to correct susceptibility distortions in spin-echo echo-planar images: application to diffusion tensor imaging. NeuroImage 20, 870–888 (2003).

8. A. Schaefer, et al., Local-Global Parcellation of the Human Cerebral Cortex from Intrinsic Functional Connectivity MRI. Cereb Cortex 28, 3095–3114 (2018).

9. B. Whitcher, V. J. Schmid, A. Thorton, Working with the DICOM and NIfTI Data Standards in R. Journal of Statistical Software 44, 1–29 (2011).

10. R Core Team, R: A language and environment for statistical computing. (R Foundation for Statistical Computing, 2020).

11. M. F. Glasser, et al., The minimal preprocessing pipelines for the Human Connectome Project. NeuroImage 80, 105–124 (2013).

12. D. Marcus, et al., Informatics and Data Mining Tools and Strategies for the Human Connectome Project. Front. Neuroinform. 5 (2011).

13. R. W. Cox, AFNI: Software for Analysis and Visualization of Functional Magnetic Resonance Neuroimages. Computers and Biomedical Research 29, 162–173 (1996).

14. M. H. A. Hendriks, N. Daniels, F. Pegado, H. P. Op de Beeck, The Effect of Spatial Smoothing on Representational Similarity in a Simple Motor Paradigm. Front. Neurol. 8 (2017).

15. M. Hanke, et al., PyMVPA: a unifying approach to the analysis of neuroscientific data. Front. Neuroinform. 3 (2009).

16. C. S. H. Oedekoven, J. L. Keidel, S. C. Berens, C. M. Bird, Reinstatement of memory representations for lifelike events over the course of a week. Scientific Reports 7, 14305 (2017).

17. D. Stawarczyk, M. A. Bezdek, J. M. Zacks, Event Representations and Predictive Processing: The Role of the Midline Default Network Core. Topics in Cognitive Science (In press) https:/doi.org/10.1111/tops.12450 (September 6, 2019).

18. J. A. Etzel, “MVPA Permutation Schemes: Permutation Testing for the Group Level.” in International Workshop on Pattern Recognition in NeuroImaging, (2015), pp. 65–68.

19. J. Schrouff, et al., PRoNTo: pattern recognition for neuroimaging toolbox. Neuroinformatics 11, 319–337 (2013).

20. D. Bates, M. Mächler, B. Bolker, S. Walker, Fitting Linear Mixed-Effects Models Using lme4. Journal of Statistical Software 67, 1–48 (2015).

21. J. Fox, S. Weisberg, An R Companion to Applied Regression, Third (Sage, 2019).

22. R. Lenth, emmeans: Estimated Marginal Means, aka Least-Squares Means. R package version 1.3.5.1. (2019).

23. T. Yanagida, misty: Miscellaneous Functions “T. Yanagida”. R package version 0.3.0. (2020).

24. D. Lüdecke, sjPlot: Data Visualization for Statistics in Social Science, R package version 2.7.1 (2019).

25. J. R. Landis, G. G. Koch, The measurement of observer agreement for categorical data. Biometrics 33, 159–174 (1977).

26. C. N. Wahlheim, J. M. Zacks, Memory guides the processing of event changes for older and younger adults. Journal of Experimental Psychology: General 148, 30–50 (2019).

27. L. Hasher, R. T. Zacks, “Working memory, comprehension, and aging: A review and a new view” in The Psychology of Learning and Motivation: Advances in Research and Theory, Vol. 22, (Academic Press, 1988), pp. 193–225.

28. D. H. Kausler, Learning and memory in normal aging (Academic Press, 1994).

29. C. N. Wahlheim, B. H. Ball, L. L. Richmond, Adult age differences in production and monitoring in dual-list free recall. Psychology and Aging 32, 338–353 (2017).

30. R. Leech, D. J. Sharp, The role of the posterior cingulate cortex in cognition and disease. Brain 137, 12–32 (2014).

31. R. Leech, R. Braga, D. J. Sharp, Echoes of the brain within the posterior cingulate cortex. J Neurosci 32, 215–22 (2012).

32. M. C. Anderson, J. H. Neely, “Interference and inhibition in memory retrieval.” in Memory, 2nd Ed., E. L. Bjork, R. A. Bjork, Eds. (Academic Press, 1996), pp. 237–313.

33. C. M. Bird, How do we remember events? Current Opinion in Behavioral Sciences 32, 120–125 (2020).

34. W. D. Spencer, N. Raz, Differential effects of aging on memory for content and context: A meta-analysis. Psychology and Aging 10, 527–539 (1995).

35. K. A. Norman, E. L. Newman, G. Detre, A neural network model of retrieval-induced forgetting. Psychological Review 114, 887–953 (2007).

36. M. Brett, J.-L. Anton, R. Valabregue, J.-B. Poline, Region of interest analysis using an SPM toolbox in (2002).

37. P. Zeidman, E. A. Maguire, Anterior hippocampus: the anatomy of perception, imagination and episodic memory. Nat Rev Neurosci 17, 173–82 (2016).

38. A. Kafkas, D. Montaldi, How do memory systems detect and respond to novelty? Neuroscience Letters 680, 60–68 (2018).

39. A. C. H. Lee, K. H. Brodersen, S. R. Rudebeck, Disentangling spatial perception and spatial memory in the hippocampus: a univariate and multivariate pattern analysis fMRI study. J Cogn Neurosci 25 (2013).

40. H. Kim, Differential neural activity in the recognition of old versus new events: an activation likelihood estimation meta-analysis. Hum Brain Mapp 34, 814–36 (2013).

